# Metabolic interplay drives population cycles in a cross-feeding microbial community

**DOI:** 10.1101/2024.10.14.618235

**Authors:** Tyler D. Ross, Christopher A. Klausmeier, Ophelia S. Venturelli

## Abstract

Population cycles are prevalent in ecosystems and play key roles in determining their functions^1,2^. While multiple mechanisms have been theoretically shown to generate population cycles^3–6^, there are limited examples of mutualisms driving self-sustained oscillations. Using an engineered microbial community that cross-feeds essential amino acids, we experimentally demonstrate cycles in strain abundance that are robust across environmental conditions. A nonlinear dynamical model that incorporates the experimentally observed cross-inhibition of amino acid production recapitulates the population cycles. The model shows that the cycles represent internally generated relaxation oscillations, which emerge when fast resource dynamics with positive feedback drive slow changes in strain abundance. Our findings highlight the critical role of resource dynamics and feedback in shaping population cycles in microbial communities and have implications for biotechnology.

## Main Text

Population cycles (regular oscillations in population size) are widespread in ecological systems, with almost 30% of natural populations showing such dynamics^1^. Many mechanisms have been proposed to explain how internally generated oscillations can emerge from nonlinear interactions between populations. Beginning with Lotka and Volterra’s foundational predator-prey model^7,8^, consumer-resource (+/-) interactions have been shown to be a common cause of population cycles^3^. Other theoretical mechanisms that can generate regular oscillations include age- and stage-structure^4^, intransitive competition^5^, and eco-evolutionary dynamics^6^. Notably absent from this list is mutualism.

Mutualism (+/+) is increasingly recognized as a pervasive and influential interaction motif shaping diverse ecosystems^9–11^. Mutualistic interactions can give rise to nonlinear phenomena such as alternative stable states^12^. However, this interaction motif has not been widely considered a driver of oscillatory dynamics. Mutualisms were first modeled using the generalized Lotka-Volterra model^13^, but this model can generate unbounded growth and instability^14^. In more recent models, the mechanisms driving mutualisms are a central focus, particularly among interacting macroscopic organisms^15^. Mutualisms are also present in microbial communities, whose interactions influence a broad range of natural phenomena from biogeochemical cycles to human health and behavior^16,17^. Cross-feeding, a mechanism of mutual resource exchange, is frequently observed in microbial communities^18–22^. Consistent with the theoretical understanding that mutualisms lead to stable equilibria, current models of cross-feeding mutualisms do not exhibit population cycles^23^.

Here we demonstrate that an engineered microbial community shaped by bidirectional cross-feeding of essential amino acids can exhibit robust population cycles across a range of environmental conditions. Resource profiling demonstrates a pattern of amino acid cross-inhibition that has not been previously captured in models of mutualisms. A nonlinear dynamical systems model that encodes the observed mutual resource inhibition replicates the observed dynamics and can accurately extrapolate to new environmental conditions. Our model demonstrates that the cycles emerge as internally generated relaxation oscillations, where the fast mutual inhibition of amino acid production (positive feedback) leads to alternative stable states, which drive slow population dynamics. Our results demonstrate that resource-mediated interactions in microbial communities can generate rich dynamical behaviors and have implications for innovations in biotechnology.

### Microbial communities driven by amino acid cross-feeding can exhibit distinct dynamical behaviors

To investigate how cross-feeding shapes community dynamics, we engineered individual *Escherichia coli* amino acid auxotrophs *ΔtyrA* and *ΔpheA* that reciprocally cross-feed phenylalanine and tyrosine respectively, while competing for glucose (**Fig. 1a**). Consistent with this mechanism, monocultures exhibited minimal growth after 24 hours in media lacking amino acids, whereas co-cultures showed substantial growth of both strains (**Fig. 1b**). This shows that cross-feeding plays a critical role in structuring communities in environments lacking external supply of the required amino acids.

**Figure 1.**
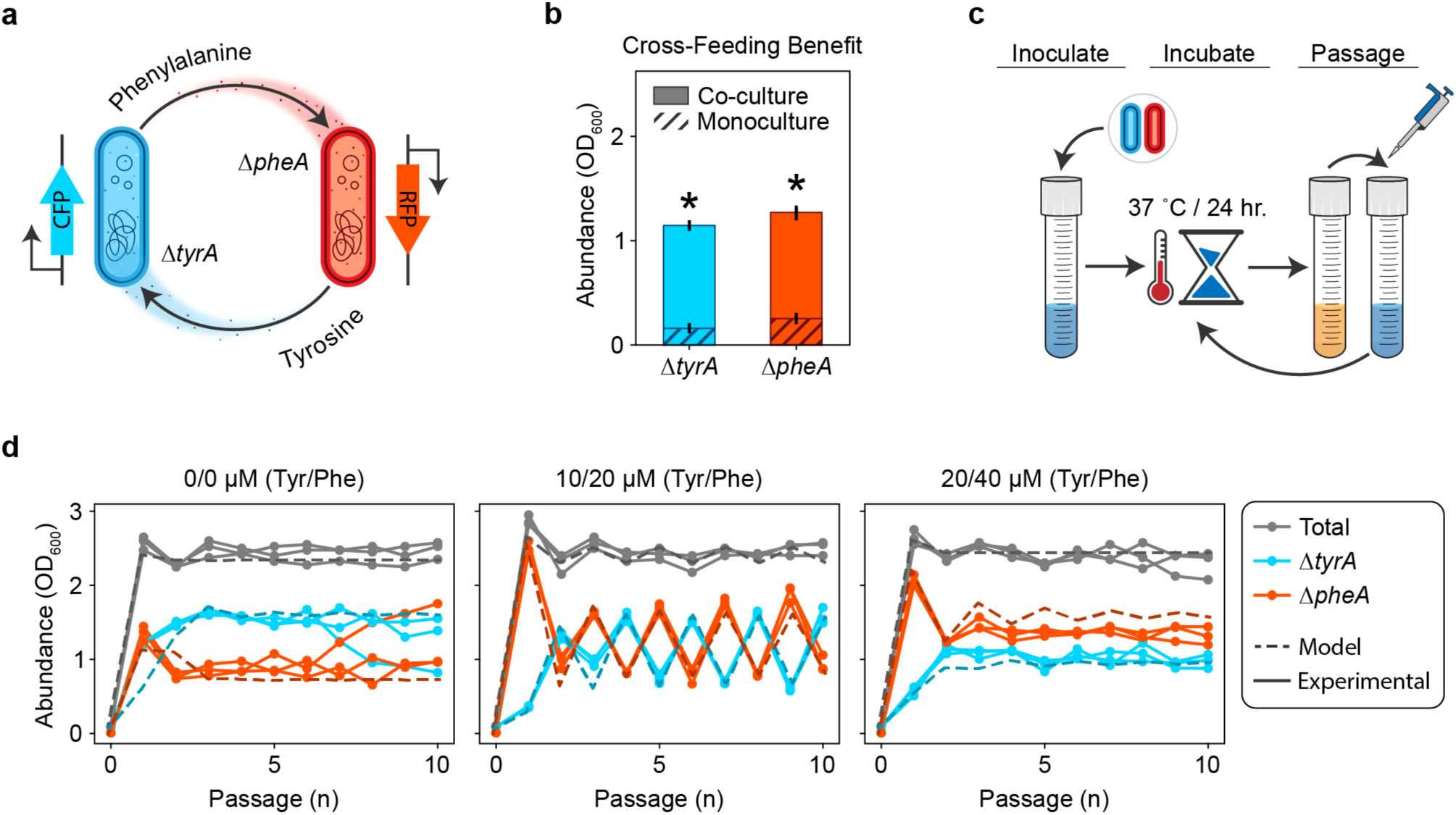
An amino acid cross-feeding microbial community displays stable equilibrium or oscillatory dynamics. **a**, Two *E. coli* amino acid auxotrophs exhibit mutually beneficial cross-feeding. Mutant *ΔtyrA* produces phenylalanine for mutant *ΔpheA*, while mutant *ΔpheA* reciprocally produces tyrosine for mutant *ΔtyrA*. Each auxotroph harbors an IPTG inducible fluorescence construct for strain identification and abundance quantitation in co-culture. **b,** Relative growth benefit due to cross-feeding. Auxotrophs *ΔtyrA* and *ΔpheA* were grown as monocultures and a pairwise community in minimal medium lacking any amino acid supplementation. The absolute abundance after 24 hours of growth for each monoculture are shown with dashed bars. The absolute abundance after 24 hours of growth for each member in the pairwise community is shown with solid bars. Asterisks indicate statistical significance between the average growth in monoculture versus co-culture (*ΔtyrA*: p = 1.5220*E* − 05, *ΔpheA*: *p* = 7.8113*E* − 06). **c,** In the experimental passaging scheme, a minimal or supplemented media is inoculated with a pair of cross-feeding amino acid auxotrophs. The culture is incubated for a period of 24 hours prior to being diluted to a constant OD_600_. Cycles of growth and dilution are carried forward for a finite number of passages. **d,** Dynamical behaviors of the *ΔtyrA* / *ΔpheA* community when subjected to the experimental passaging scheme shown in (**c**). Separate plots correspond with different concentrations of amino acid supplementation in the passaging media. Grey lines indicate the abundance of the full community while colored lines indicate the absolute abundance of *ΔtyrA* (cyan) or *ΔpheA* (red). Solid lines with circular markers represent experimental data and dashed lines represent simulations of equation (1).

To investigate the dynamics of the community, we co-cultured the strains in serial batches subjected to daily dilution with fresh media for ten days, with no, low and moderate levels of external amino acid supply (**Fig. 1c**). With no or moderate external amino acid supply, the composition of the communities converged to an equilibrium (**Fig. 1d**). Notably, the community exhibited sustained large-amplitude period-two oscillations with low concentrations of external amino acids. To evaluate the robustness of these dynamics to environmental parameters, we varied the external amino acid supply over a broader range of concentrations (**Extended Data Fig. 1**). The community displayed period-two oscillations across a range of intermediate amino acid supply concentrations. These results demonstrate that oscillatory dynamics can emerge in microbial communities whose dominant interaction is resource cross-feeding. In addition to oscillations and stable coexistence, *ΔtyrA* was excluded when the external phenylalanine supply was high and tyrosine supply was low, indicating that the contribution of mutualism versus resource competition was context dependent.

### Tyrosine and phenylalanine release primarily occurs during focal amino acid starvation

Extant models of cross feeding that do not produce oscillations assume that nutrients are released at constant rates^23,24^. The presence of population cycles in our cross-feeding community suggests that these models neglect additional feedback loops required for generating oscillations. In accordance with previous results that demonstrate a time-dependent change in the release of certain metabolites^25^, we hypothesized that there exists unresolved feedback shaping the dynamics of amino acid release rates. Therefore, we measured the concentrations of extracellular resources (amino acids and glucose) over time for each auxotroph in response to varying supplementation of the required amino acid (**Fig. 2a**). When high concentrations of the required amino acid were provided, a residual amount remained at the onset of stationary phase as glucose was depleted (**Fig. 2b,e**). In these cases of glucose limitation, almost none of the partner’s amino acid was released at 12 hours, and only a small amount was released at 24 hours, likely due to cell death and lysis^26^. Conversely, the required amino acid was completely depleted in stationary phase when lower amino acid concentrations were initially supplied. A moderate amount of remaining glucose indicated that these cultures were amino acid, as opposed to glucose, limited (with some exception at 24 hours, see Supplementary Information). Notably, for both *ΔtyrA* and *ΔpheA*, the limitation of required amino acid was associated with a substantial release of the partner strain’s required amino acid (**Fig 2c,d,f,g**). These results demonstrate reciprocal inhibition of amino acid release, where tyrosine inhibits release of phenylalanine by *ΔtyrA* and vice versa. Such cross-inhibitory interactions generate a positive feedback loop in the resource dynamics. When a given auxotroph is limited by its required amino acid, it releases the partner’s required amino acid (producer). This leads to glucose limitation of the partner strain as opposed to amino acid limitation, preventing the reciprocal release of amino acid (consumer). Amino acid limitation of the producer is thus reinforced through a net positive feedback loop due to the cross-inhibition topology.

**Figure 2.**
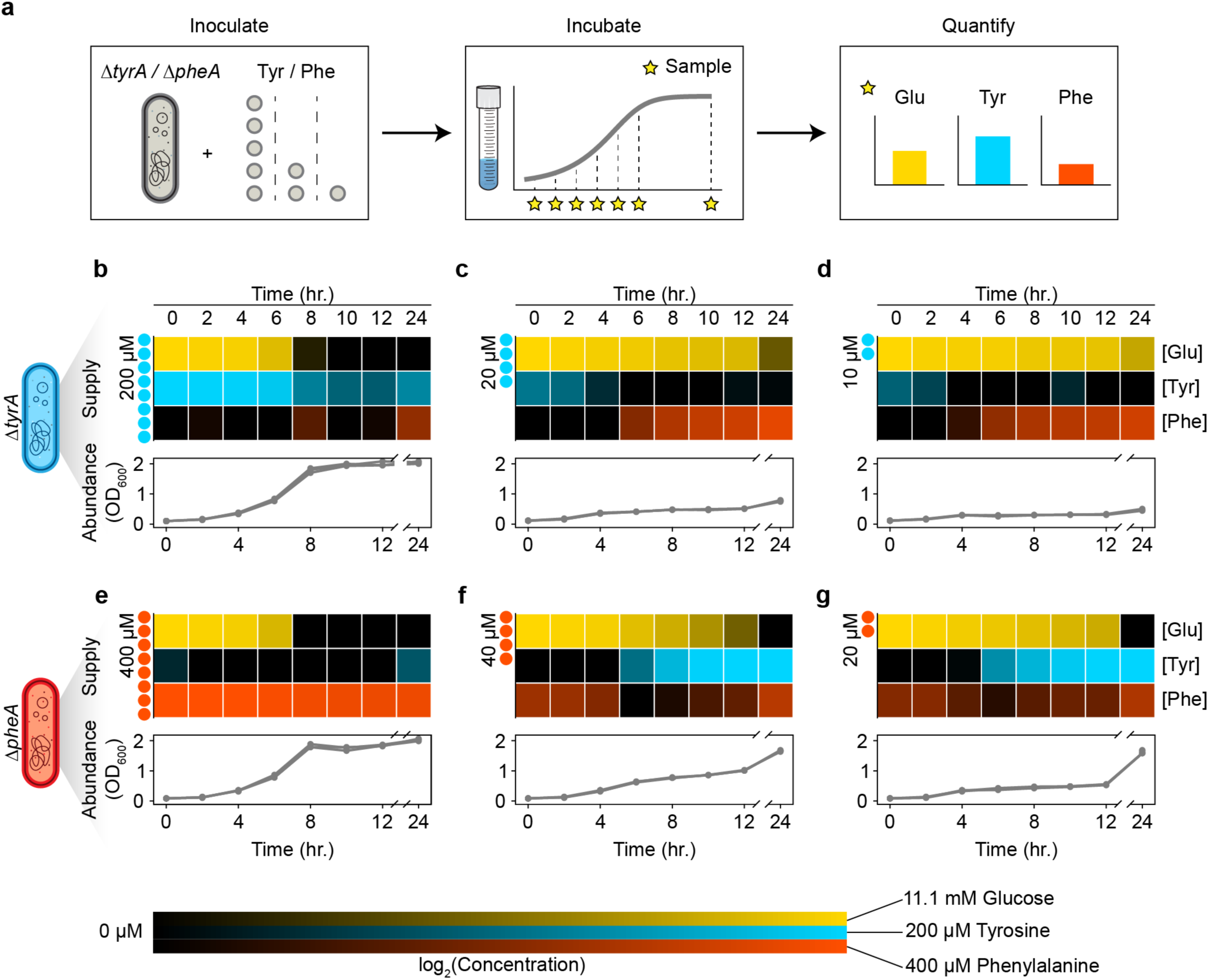
Extracellular metabolite measurements demonstrate that amino acid release occurs in response to amino acid limitation. **a**, Experimental setup to analyze the extracellular dynamics of essential auxotroph resources. Each auxotroph was cultured as a monoculture in media with either 200 µM, 20 µM, or 10 µM tyrosine for *ΔtyrA*, or 400 µM, 40 µM, or 20 µM phenylalanine for *ΔpheA*. Cultures were sampled every 2 hours over a 12-hour period, and again at 24 hours. Glucose, tyrosine, and phenylalanine concentrations were quantified using fluorometric and colorimetric assay kits. **b-g,** Extracellular resource dynamics in auxotroph monocultures. Line plots represent growth curves at different concentrations of amino acid supplementation, while heatmaps depict corresponding extracellular resource concentrations. Panels (**b-d**) show *ΔtyrA* monocultures, and panels (**e-g**) show *ΔpheA* monocultures. The initial amino acid supplementation levels (tyrosine for *ΔtyrA* and phenylalanine for *ΔpheA*) are indicated to the left of each heatmap. Heatmap shading indicates the concentrations of glucose (yellow), tyrosine (cyan), and phenylalanine (red).

### A dynamic model capturing resource and strain dynamics recapitulates system behaviors

To determine if the cross-inhibition of amino acid release can explain the emergence of oscillatory dynamics in our engineered community, we constructed a nonlinear ordinary differential equation model of the system. Our model considers two auxotrophs (*ΔtyrA* and *ΔpheA*, denoted *N*_1_ and *N*_2_, equations (1a) and (1b)), two cross-fed resources (tyrosine and phenylalanine, denoted *R*_1_ and *R*_2_, equations (1c) and (1d)) and one resource that ultimately limits the community (glucose, denoted *R*_3_, equation (1e))^23^. Each auxotroph *i* has two essential resources (its required amino acid and glucose), with its realized growth rate μ_i_ given by the minimum of its potential growth on the two resources *j*, μ_ij_(*R*_j_), which take Michaelis-Menten form (equations (1f) and (1g)). Amino acid uptake is proportional to the realized growth rate of the auxotrophs, while glucose uptake is proportional to their glucose-limited growth rate, with stoichiometric coefficients *q*_ij_. We assume auxotrophs produce amino acid to match their glucose uptake and release any excess to the environment at rate *q*_ii_(μ_i3_ – μ_i_). When growth is glucose-limited, μ_i_ = μ_i3_, and amino acids are not released into the environment. This mass-balance constraint results in amino acid release only when glucose is available and the required amino acid limits growth, consistent with our experimental results (**Fig. 2**).

We model the external supply of resources with concentration *R*_j,in_ and dilution of species and resources at rate *D*. For comparison with our experimental results, we initially treat the dilution rate as a series of discrete events using *D*(*t*) = *f* ∑_*n*=1_ δ(*t* − *n*τ), where *f* is the fraction of culture transferred, *n* is the passage number, τ is the period, and δ(⋅) is the Dirac delta function. For further theoretical exploration, we treat the dilution rate *D* as a constant, as in a chemostat. Together, these assumptions result in the following model:

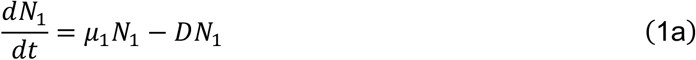

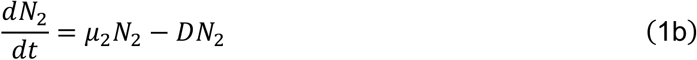

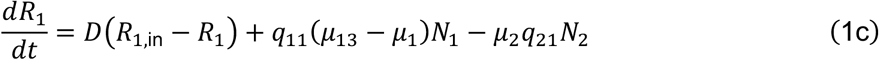

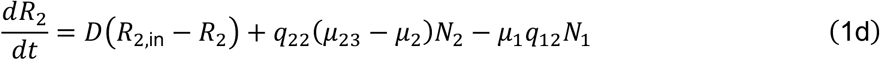

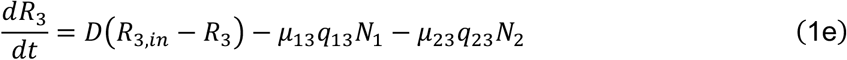

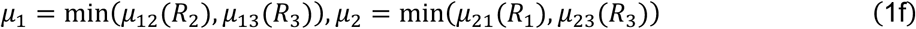

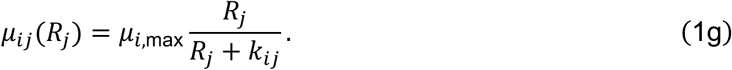

A complete description of the model parameters is provided in Extended Data Table 1.

We inferred the model parameters using measurements of absolute abundance from our initial ten-batch passaging experiment (see Methods). Our model and best estimate parameter set accurately fit the different community dynamical behaviors as a function of the amino acid supply concentration (**Fig. 1d**). Notably, the model accurately predicted the dynamical behaviors for various amino acid supply concentrations that were not used for model fitting as well as the batch growth dynamics of a cycling community (Extended Data Fig. 1 and 2). This demonstrates that the model could extrapolate to new environments. These results suggest that cross-feeding with reciprocal inhibition can generate the period-two cycles observed experimentally. We also explored an alternative model that uses predicted regulatory links at the biomolecular level to encode a more explicit mechanism of resource cross inhibition^27–30^. This alternative model could recapitulate the period-two cycles and equilibrium dynamics as a function of the amino acid supply concentrations (see Supplementary Information). This implies that the resource cross inhibition topology was critical for generating oscillations in the system.

Cycles can emerge due to external forcing or be self-sustained due to internal mechanisms driving the dynamics^31^. For example, periodic forcing through daily passaging was necessary for the development of cycles in a mutualistic cross-protection microbial community^32^. To determine whether cycles can emerge in the absence of external forcing, we analyzed our model with constant dilution rate *D* as in a chemostat (**Fig. 3a**). Our results show that periodic limit-cycle solutions occur across a range of low external amino acid supply (**Fig. 3b**). This suggests that the experimentally observed cycles are not dependent on periodic forcing through daily passaging but arise due to the internal dynamics of the community. Higher amino acid supply concentrations lead to stable equilibria, with a substantial range of parameters leading to stable coexistence of the auxotrophs. However, competitive exclusion occurs at extreme supply concentrations, and the outcome of this competition is solely sensitive to tyrosine supply as opposed to the tyrosine-to-phenylalanine supply ratio^33^. This reflects the *ΔtyrA* auxotroph’s superior competitive ability for glucose using the inferred parameter set, which becomes limiting when sufficient tyrosine is externally supplied. Notably, this pattern of sensitivity can switch depending on moderate parameter variations dictating the glucose utilization efficiency of each strain (**Extended Data Fig. 3**). This implies that the pattern of tyrosine supply sensitivity is not robust to variations in these parameters.

**Figure 3.**
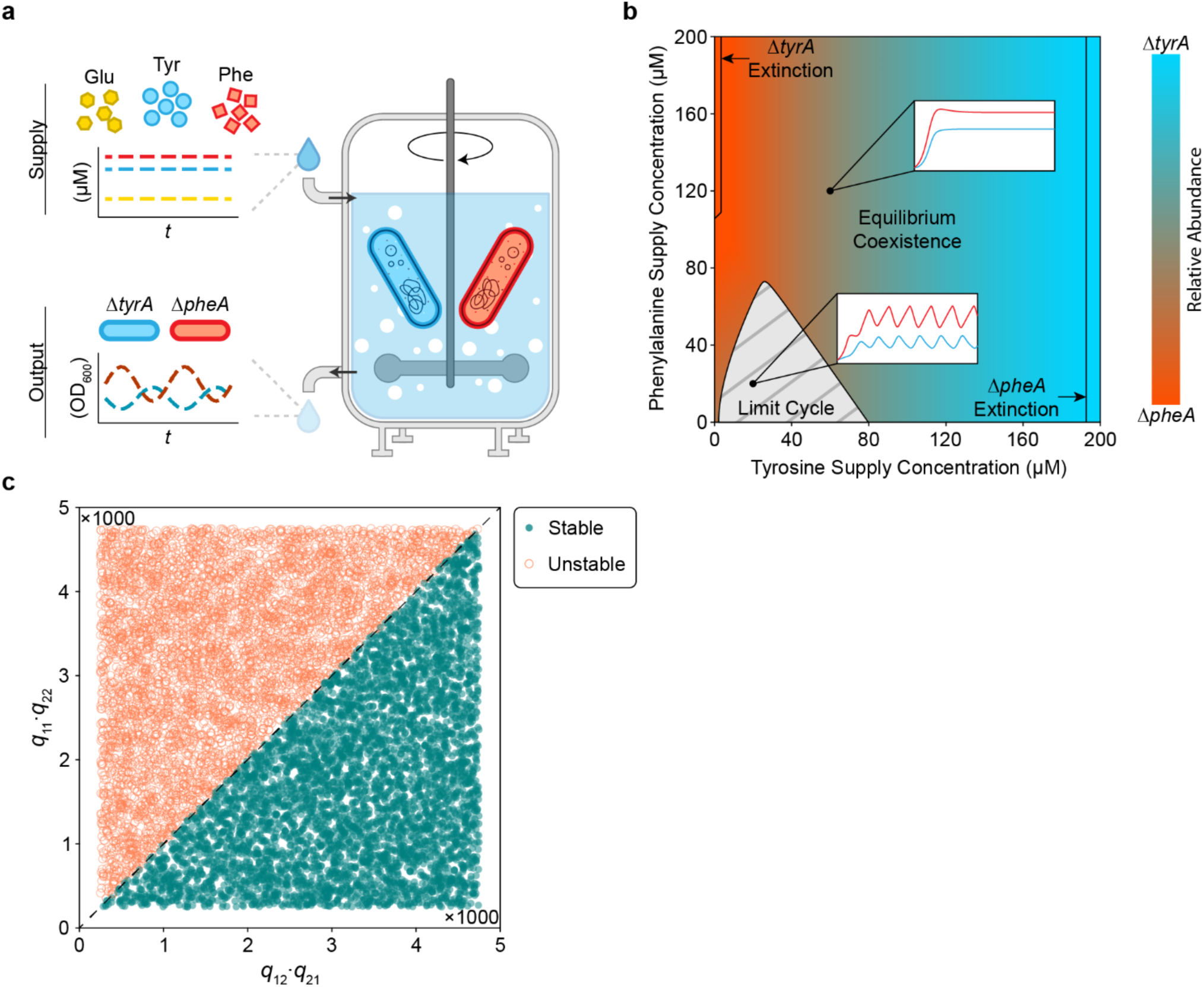
Population cycles exist in the absence of external forcing. **a**, Schematic representation of the *in silico* chemostat experiment. The chemostat facilitates continuous growth of the *ΔtyrA* / *ΔpheA* community through constant inflow of glucose, tyrosine, and phenylalanine. Perfusion is governed by the dilution rate *D*, which acts on the inflow of external resource, and the outflow of chemostat resource and excess cells. **b,** Two-dimensional bifurcation diagram of the *ΔtyrA* / *ΔpheA* community dynamics in a virtual chemostat. As the amino acid supply concentrations vary along each axis, the steady state community composition is indicated according to the color bar. A grey hashed region indicates the amino acid supply concentrations that do not produce stable equilibria, but instead display limit cycle oscillations. Transcritical bifurcations beyond which the community collapses into a monoculture are indicated with solid black lines. Two inset plots demonstrate simulations of stable equilibrium and cycling dynamics. **c,** Steady state stability for randomly sampled values of *q*_ij_. Jacobian matrix stability analysis was used to determine the steady state stability for a range of *q*_11_*q*_22_and *q*_12_*q*_21_from 250 to 4750. Each point in the scatter plot represents the stability of a steady state for a given parameter sample. Filled in blue points represent stable steady states while empty orange circles represent unstable steady states. A black line separates the region where *q*_11_*q*_22_ > *q*_12_*q*_21_.

### Dissecting the mechanisms that drive cycles

To more deeply understand the mechanism driving the cyclic population dynamics, we derived a minimal model for the two amino acids (*R*_1_ and *R*_2_) and the relative abundance of the two auxotrophs, *f* = *N*_1_/(*N*_1_ + *N*_2_) (see Supplementary Information for a full derivation and additional discussion). This minimal model captures the essential features of our system, shows similar dynamics to equation (1) (**Extended Data Fig. 4**), and is more amenable to analytical methods. This analysis identifies the cycles as relaxation oscillations, which show fast-slow dynamics^34,35^, and have been identified in some predator-prey models^36^. Based on the parameter values inferred above, resource dynamics occur on a fast timescale and can show two alternative stable metabolic states if the positive feedback due to resource cross-inhibition is stronger than negative feedback due to self-damping (growth-imposed resource limitation). In each alternative state, one consumer auxotroph is glucose-limited, so it does not release amino acids. The other, producer auxotroph is consequently limited by its required amino acid and therefore releases the amino acid it produces, which fuels growth of the consumer. This matches the experimentally observed intra-batch resource dynamics (**Fig. 2**). The asymmetric cross-feeding relationship between the auxotrophs drives the strain relative abundance dynamics on a slower timescale. As the relative abundance of the producer auxotroph decreases, the amino acid concentrations track the stable resource equilibrium until it vanishes at a saddle-node bifurcation (**Extended Data Fig. 5**). This triggers an abrupt jump to the alternative resource equilibrium where the roles of producer and consumer auxotroph are reversed. Simulation of the full model (equation (1)) shows similar dynamics (**Fig. 4**) except for a brief period where both amino acids are being produced that coincides with the switch between alternative metabolic states (**Fig. 4b** iii and vi).

**Figure 4.**
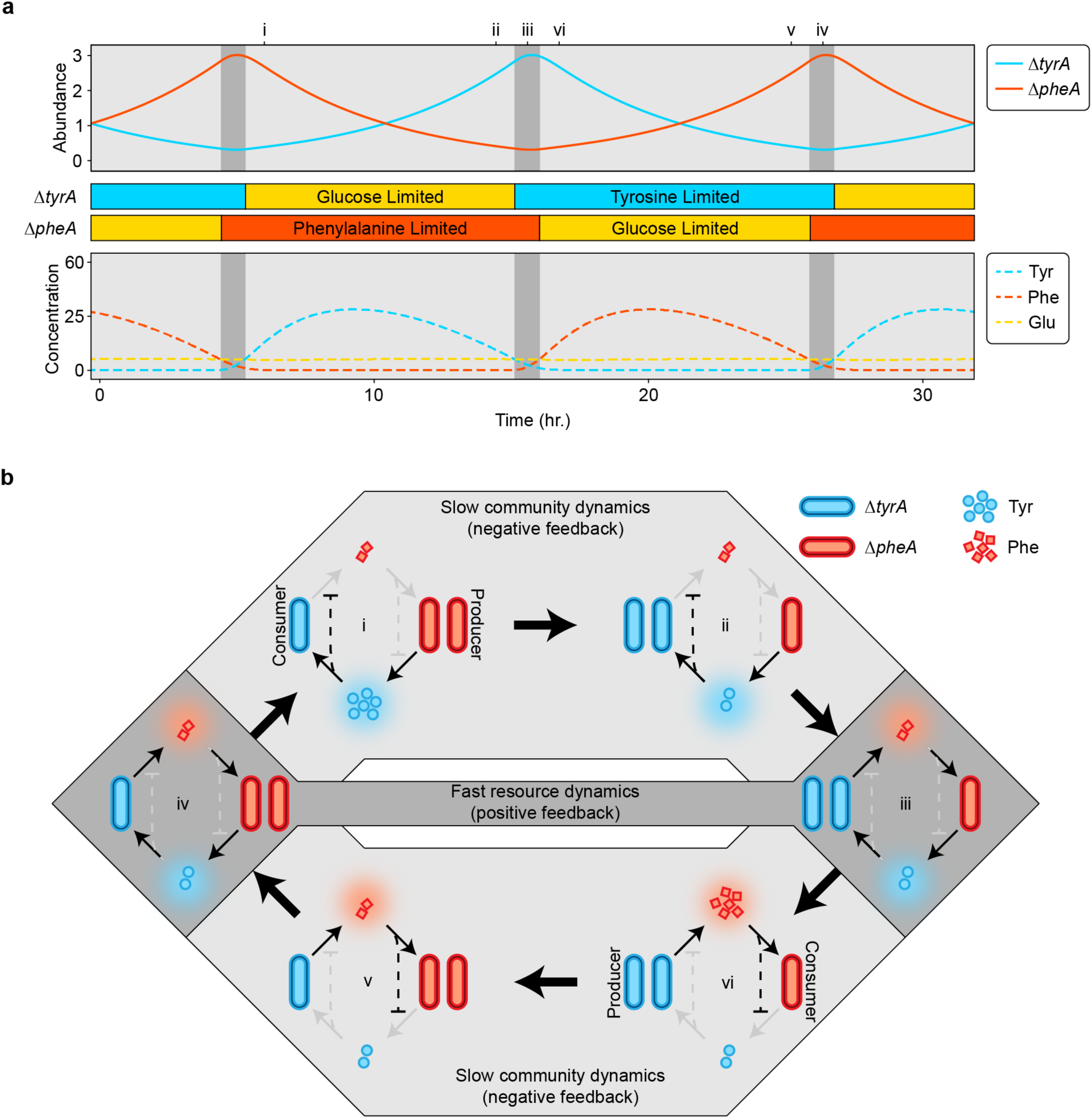
Continuous time simulations highlight the interplay of slow community dynamics and fast resource dynamics in a cross-feeding system. **a**, Simulations of auxotroph abundance (top) and resource concentration (bottom) are shown for a set of chemostat parameters where *D* = 0.2, *R*_1,i*n*_ = 1, *R*_2,i*n*_ = 1, and *R*_3,i*n*_ = 11.11. The auxotroph parameters were simplified and made to be symmetrical for illustrative purposes (Extended Data Table 1). Colored bars between the plots of abundance and resource dynamics show the intervals of glucose and amino acid limitation for each auxotroph. Shading within the plots highlights distinct types of resource limitation and interaction topography. In the light grey regions, one auxotroph is amino acid limited while the other is glucose limited. In the dark grey regions, both auxotrophs are amino acid limited. **b,** Population cycles and associated changes in the community interaction network. Each network diagram represents a snapshot of the system at the time indicated in (**a**) with a numeral (i-iv). Solid lines with arrowheads represent positive interactions, i.e., amino acid promoting the growth of either auxotroph. Dashed lines with flatheads represent inhibitory interactions. Coloring of the interaction lines corresponds with active (black) or inactive (grey) interactions. Shading around the interaction networks corresponds with the shading from (**a**), where light grey network diagrams represent slow community dynamics and dark grey diagrams represent fast resource dynamics.

The minimal model provides insights into the parameter conditions that produce cycles. Linear stability analysis shows that *q*_11_*q*_22_ > *q*_12_*q*_21_ is a necessary but not sufficient condition for existence of the alternative stable states in the fast resource-subsystem that are required for relaxation oscillations (see Supplementary Information). The parameters *q*_11_and *q*_22_represent the strength of the positive feedback loop generated by resource cross-inhibition, while *q*_12_ and *q*_21_ represent the strength of the negative feedback due to self-damping. To evaluate whether this condition can predict the emergence of oscillations in the full model (equation (1)), we randomly sampled values for *q*_ij_and determined the qualitative behavior of the system for each parameter set (**Fig. 3c**). Our results from the full model were consistent with the condition derived from the minimal model. In addition, increasing the chemostat dilution rate *D* contributes to self-damping of the resources (i.e. negative feedback), which reduces the likelihood of alternative stable steady states and therefore cycles. In sum, our findings highlight the critical role that resource feedback and dynamics play in shaping the complex behaviors of microbial communities.

## Discussion

Previous models of mutualisms in general, and cross-feeding in particular, do not predict self-sustained oscillations in the absence of external forcing. The salient difference in our system is the feedback generated by amino acid release in response to amino acid limitation of the producer auxotroph (**Fig. 2b**). This represents a strategy of resource management involving the release of excess produced amino acid during periods of limitation by the required amino acid. This strategy could be explained by an endogenous stress response mechanism in *E. coli* in which specific amino acids, including the aromatics, accumulate during starvation because of protein degradation ^25,37,38^. As an alternative explanation, the observed resource dynamics could be an indirect consequence of the auxotrophic gene knockouts. Metabolic flux intended to remedy starvation of the required amino acid could easily leak into an overlapping pathway (see Supplementary Information). Indeed, the anabolic pathways for tyrosine and phenylalanine are nearly identical with only the final two reactions being distinct, suggesting that dysregulation of one pathway could impact the metabolic fluxes of the other pathway. Beyond the specific mechanism, the resulting positive feedback loop captured in our models is essential to the oscillations in our system. Further, similar features of our system may occur more broadly in ecological systems shaped by resource exchange beyond microbial communities. A plant growth model with separate root and shoot compartments that share excess resources (nitrogen and carbon) shows similar dynamics, alternating between two metabolic equilibria^39,40^.

Although the interaction between strains is mutualistic (+/+) when averaged over an entire cycle, at most instants one auxotroph benefits without concurrently returning the favor, representing a transient commensalism. Due to the oscillations, species take turns as they alternate out of phase between disjoint periods of growth and amino acid resource production (**Fig. 4a**). Such dynamics mirror reciprocal altruism, where each strain temporarily sacrifices its growth to benefit the other, ensuring long-term mutual benefit^41^.

One consequence of these oscillations is that they can prevent the disruption of the mutualism by a cheater that consumes the exchanged resources without producing either of these resources (e.g. a dual auxotroph) in model simulations. In conditions where the community converges to a stable equilibrium, such a cheater can invade a community of two cross-feeding auxotrophs that are each deficient in producing a single amino acid. In this case, the community collapses as the mutualists go extinct in response to depletion of the exchanged resources by the cheater^23^. Because the cheater requires both exchanged resources, any mechanism that separates their simultaneous availability can prevent a successful invasion. Previously, we have shown that local interactions between cross-feeding auxotrophs can exclude cheaters by spatially separating the exchanged resources^42^. Notably, temporal separation of resources, due to the alternating production of individual amino acids in the oscillating community (**Fig. 4a**), provides another mechanism to exclude an otherwise successful cheater (**Extended Data Fig. 6**). This implies that oscillations may represent an ecological strategy to resist invasion by cheaters, which could be leveraged for engineering stable microbial communities for biotechnology applications.

## Methods

### Strains

Strains used for all experiments were *E. coli* K-12 BW25113 single amino acid auxotrophs derived from the Keio Knockout Collection^43^. Deletion of the *tyrA* and *pheA* genes established the tyrosine and phenylalanine auxotrophies respectively. Each auxotroph was transformed with a pBbA6c plasmid^44^ harboring a chloramphenicol resistance gene for plasmid maintenance, and an isopropyl β-D-1-thiogalactopyranoside (IPTG) inducible fluorescence marker. The auxotrophs were transformed such that *ΔtyrA* expressed CFP and *ΔpheA* expressed RFP.

### Preculture conditions

For each experiment involving the *ΔtyrA* or *ΔpheA* auxotrophs, precultures were prepared by inoculating EZ Rich Defined Media (Teknova, #M2105) (Extended Data Table 3) containing 25 µg/mL chloramphenicol with approximately 2 µL of the appropriate glycerol stock. Cultures were then incubated for 16 hours at 37 °C with 250 RPM orbital shaking. Cultures were then centrifuged for 5 minutes at 3000 RCF, and the cell pellet was washed once with MOPS Minimal Media (Teknova, #M2106) prior to resuspension.

### Microscopy and cell counting

Absolute abundance measurements of the auxotrophs in co-culture were determined by combining OD_600_ reading of the co-culture with relative abundance measurements. Relative abundance was measured using epifluorescence microscopy and a cell counting script that categorizes and counts cells based on their fluorescent label. For each culture, a 2 µL sample was spotted onto a glass slide treated with poly-L-lysine (Sigma, #P8920) and covered with a glass coverslip. Microscopy was performed on a Nikon Eclipse Ti where two regions of interest were imaged for each sample using a 20X objective. Each image was composed of a phase contrast channel, an ECFP channel for *ΔtyrA* identification (Nikon, #96361, 436×20 nm excitation / 480×40 nm emission), and an mCherry channel for *ΔpheA* identification (Nikon, #96365, 560×40 nm excitation / 630×70 nm emission).

The total number of cells in each fluorescence channel was counted using an automated script in ImageJ. The script applies a Gaussian blur filter (σ = 1) prior to employing the find maxima function for individual cell identification. A peak prominence value of 300 was used for the find maxima function and cells on the edge of the image were excluded from the final count. The relative abundance of each species was calculated by dividing the number of cells in either fluorescence channel by the total number of cells in both fluorescence channels.

### Obligate exchange experiment (*Fig. 1b*)

Precultures of *ΔtyrA* and *ΔpheA* were diluted into 1 mL of fresh MOPS Minimal Media (Teknova #M2106) (Extended Data Table 3) containing 25 µg/mL chloramphenicol and 1 mM IPTG as monocultures and a pairwise coculture. Each culture was inoculated to achieve a total optical density at 600 nanometers (OD_600_) of 0.1. For the coculture, the inoculum ratio of the two auxotrophs was 1:1. All cultures were then incubated for 24 hours at 37 °C with 250 RPM orbital shaking. Following incubation, the OD_600_ of each culture was measured. Relative abundance was also measured in the community culture using epifluorescence microscopy (Nikon Eclipse Ti) and an automated cell counting script (ImageJ). Absolute abundance of each strain was calculated by multiplying the relative abundance measurements with the OD_600_ of the community.

### Batch passaging experiments (*Fig. 1d* and Extended Data Fig. 1)

Stock solutions of L-tyrosine (Sigma-Aldrich, #T3754) and L-phenylalanine (Dot Scientific, #DSP20260) were prepared in Milli-Q water, and subsequently used to prepare supplemented MOPS Minimal Media. The tyrosine stock solution was prepared at a 1 mM concentration in warm water with 1 hour of sonication to facilitate dissolution. The phenylalanine stock solution was prepared at a 40 mM concentration with stirring. For the initial batch passaging experiment (Fig. 1d), the amino acid supply concentrations were either 0 µM, 10 µM, or 20 µM for tyrosine, and 0 µM, 10 µM, or 20 µM for phenylalanine. For the expanded batch passaging experiment (Extended Data Fig. 1), the supply concentrations of tyrosine were 0 µM, 5 µM, 10 µM, 20 µM, 40 µM, or 80 µM, while the concentrations of phenylalanine were 0 µM, 10 µM, 20 µM, 40 µM, 80 µM, or 160 µM.

For all batch passaging experiments, precultures of *ΔtyrA* and *ΔpheA* were diluted into 1 mL of fresh media (n=3) containing 25 µg/mL chloramphenicol and 1 mM IPTG as pairwise communities. The total OD_600_ of the inoculum was equal to 0.1 for all conditions. Communities were established at a ratio of 9:1 (*ΔtyrA* to *ΔpheA*). This inoculum bias was introduced to synchronize the oscillation phase across replicates and was not necessary for the development of oscillations (Extended Data Fig. 7). Cultures were then incubated at 37 °C with 250 RPM orbital shaking and passaged every 24 hours to an OD_600_ of 0.1 in fresh media. At each passage, the relative abundance of each auxotroph was measured using epifluorescence microscopy (Nikon Eclipse Ti) and an automated cell counting script (ImageJ).

### Amino acid and glucose profiling experiment (*Fig. 2*)

Precultures of *ΔtyrA* and *ΔpheA* were diluted as monocultures (n=3) into 4 mL of fresh MOPS Minimal Media (Teknova, #M2106) containing 25 µg/mL of chloramphenicol. Monocultures of *ΔtyrA* were supplemented with either 10 µM, 20 µM, or 200 µM of tyrosine, and *ΔpheA* was supplemented with either 20 µM, 40 µM, or 400 µM of phenylalanine. Following inoculation, the cultures were incubated with shaking for 24 hours (37 °C / 250 RPM). Every two hours during the first 12 hours of incubation, the OD_600_ of each culture was measured using a NanoDrop spectrophotometer (Thermo, #ND-ONEC-W) and 300 µL samples were extracted for further processing.

Purified conditioned media was obtained from each sample through centrifugation and aspiration of the supernatant, followed by filtration using a 0.2 µM hydrophilic polyethersulfone membrane (Pall, #8019). A final set of samples were obtained at 24 hours of incubation.

Quantitation of tyrosine, phenylalanine, and glucose in the purified conditioned media samples was achieved with separate enzymatic assay kits and a multimode microplate reader (Tecan Spark). Tyrosine was quantified with a colorimetric kit (Cell Biolabs, #MET5073), phenylalanine was quantified with a fluorometric kit (bioAssay Systems, #EPHE100), and glucose was quantified with a fluorometric kit (Invitrogen, #A22189).

### Intrabatch dynamics experiment (Extended Data Fig. 2)

Precultures of *ΔtyrA* and *ΔpheA* were used to inoculate 1 mL of fresh MOPS Minimal Media (Teknova, # M2106) supplemented with 10 µM tyrosine, 20 µM phenylalanine, 1 mM IPTG, and 25 µg/mL chloramphenicol with three replicates for each condition. The inoculum ratio of *ΔtyrA* to *ΔpheA* was 1:9 and the total community OD_600_ was equal to 0.1. Cultures were then incubated at 37 °C with 250 RPM orbital shaking and passaged every 24 hours by diluting the culture to an OD_600_ of 0.1 at the beginning of each passage. At each passage, the community composition was analyzed using epifluorescence microscopy (Nikon Eclipse Ti) and an automated cell counting script (ImageJ). Daily passages into fresh media and measurements of auxotroph abundance proceeded for a total of six days.

During the fifth and sixth batches, samples of each culture were taken every two hours for the initial eight hours, then once more at 24 hours. Sampling within the batch was performed without passaging, and steps were taken to minimize the duration that cultures were outside of the incubator. Each sample consisted of 10 µL, of which 2 µL was used for relative abundance quantification with epifluorescence microscopy, and another 2 µL was diluted to 20 µL for quantification of OD_600_ with a NanoDrop spectrophotometer (Thermo, #ND-ONEC-W).

### Parameter inference

The differential equation model (equation (1)) was fit to the experimental data shown in Fig. 1d using the *`fmincon`* function in MATLAB (R2022a), which minimizes a constrained nonlinear multivariable objective function. We constructed an objective function that simulates the passaging experiment using *`ode89`* and predicts auxotroph abundance values corresponding with the data presented in Fig. 1d. The L2-norm of the residuals, defined as the difference between the predicted and measured abundance values, was added to a regularization error as the net output from the objective function. The regularization error was computed from the L1-norm of the parameter set. A best estimate weighting scheme was applied to the parameter values during regularization to account for the differences in parameter value magnitudes. This scheme normalizes each parameter according to its weight prior to computing the L1-norm. Parameter weights were derived from the average parameter value taken over a series of manually selected parameter sets that produced qualitatively accurate results. Finally, the L1-norm was multiplied by a regularization parameter (λ) to scale the regularization error.

Due to the sensitivity of the optimization to the initial conditions, we independently varied the regularization parameter and the initial parameter guess. The initial parameter sets were constructed from a Latin hypercube sampling design where the bounds were determined from a manually selected set of parameter sets. All parameters were assigned a non-negative constraint, while μ_1,*max*_ and μ_2,*max*_ were also assigned an upper bound constraint.

Once the parameter optimizations were complete for all regularization parameters and all initial parameter sets, the optimized parameter sets were evaluated against a validation dataset (Extended Data Fig. 1), and the final parameter set was selected based on the lowest overall mean squared error across the training and validation datasets.

### Bifurcation analysis

Bifurcation analysis was performed using the inferred parameter sets (Extended Data Table 1 and 2). Direct model simulations were performed to obtain the steady state community compositions for each set of amino acid supplementation concentrations. Hopf, fold and transcritical bifurcations were identified and followed using numerical continuation in MatCont^45^ using the default integration parameters.

## Data availability

Experimental data will be publicly available at the time of publication and available to reviewers upon request.

## Code availability

Code for fitting the dynamical models and analyzing system stability using Monte Carlo methods will be provided on GitHub (https://github.com/VenturelliLab) at the time of publication.

## Acknowledgements

We thank Yili Qian for his early insight regarding modeling approaches and strategies for parameter inference. This research was supported by the Department of Defense (W911NF-19-1-0269 to O.S.V.), the National Institutes of Health (R35GM124774 to O.S.V.), and the National Science Foundation (EF-2124800 to C.A.K.). The funding organizations were not involved in the design of the study, the collection and analysis of data, the decision to publish, or the preparation of the manuscript.

## Author contributions

T.D.R. and O.S.V. designed the study; T.D.R. performed the experiments; T.D.R., and O.S.V. analyzed the data; T.D.R. and C.A.K. developed the models; all authors analyzed the models; all authors wrote the manuscript; O.S.V. supervised the research and secured funding.

## Competing interests

The authors declare that they have no competing interests.

## Additional Information

## Supplementary Information

Supplementary Information is available for this paper.

Correspondence and requests for materials should be addressed to Ophelia Venturelli or Christopher Klausmeier.

Peer review Information (Pending)

**Reprints and permissions information** is available at www.nature.com/reprints.

---

**Extended Data Fig 1.**
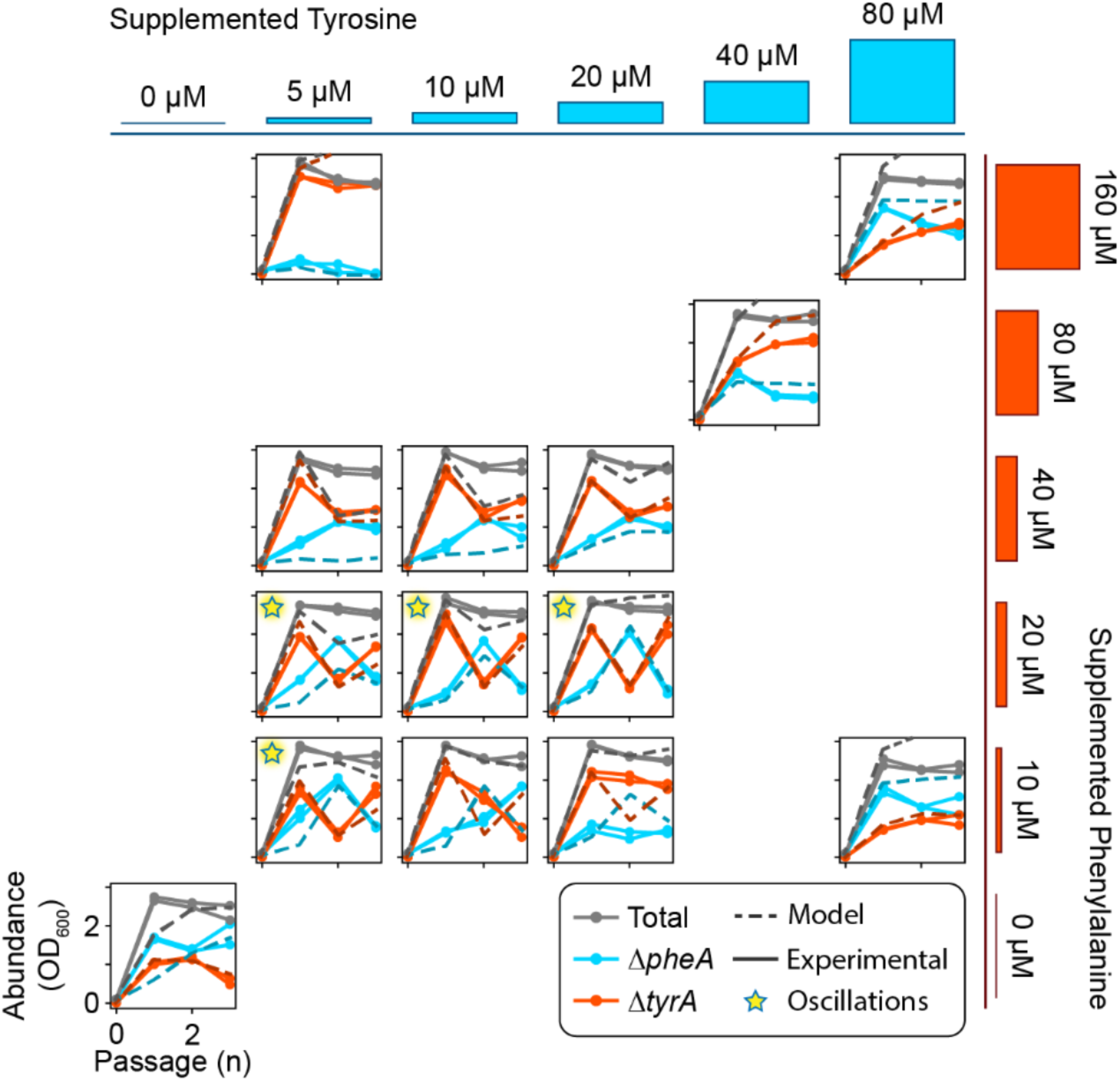
Community dynamics in response to varying concentrations of supplemented amino acids. Communities composed of *ΔtyrA* and *ΔpheA* were subjected to daily passages for three days according to the experimental scheme shown in Fig. 1c. The media was supplemented with tyrosine and phenylalanine, and the magnitude of supplementation was varied independently for each amino acid. Each subplot demonstrates the community dynamics for a distinct media composition, which is represented by the cyan and red bars on the top and right of the subplots. Cyan trajectories over time represent the absolute abundance of the *ΔtyrA* auxotroph while orange trajectories over time represent the abundance of *ΔpheA*. Solid lines correspond with experimental data while dashed lines show predictions based on the full model (equation (1)) with the inferred parameter set (Extended Data Table 1). Stars in the top left of certain subplots indicate oscillatory dynamics.

**Extended Data Fig. 2.**
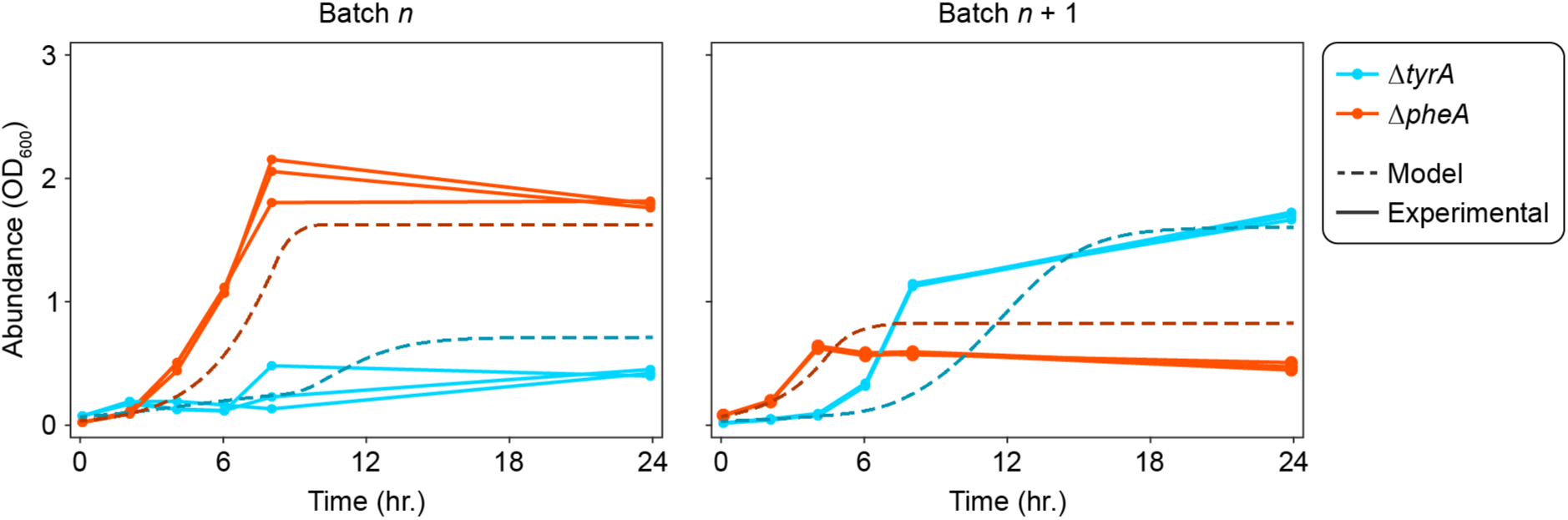
Prediction of the intra-batch dynamics for an oscillating community using the full model. Co-cultures of *ΔtyrA* and *ΔpheA* (n=3) were passaged every 24 hours in media with 5 µM tyrosine and 10 µM phenylalanine. After oscillations converged to the limit cycle (4 passages), samples were taken every 2 hours for the first 8 hours, then once more after 24 hours. The absolute abundance of each strain was measured for each sample, which is plotted as a solid line. Dashed lines represent the model prediction of the full model (equation (1)) using the inferred parameter set (Extended Data Table 1) for an initial batch (batch *n*) and a subsequent batch (batch *n* + 1).

**Extended Data Fig. 3.**
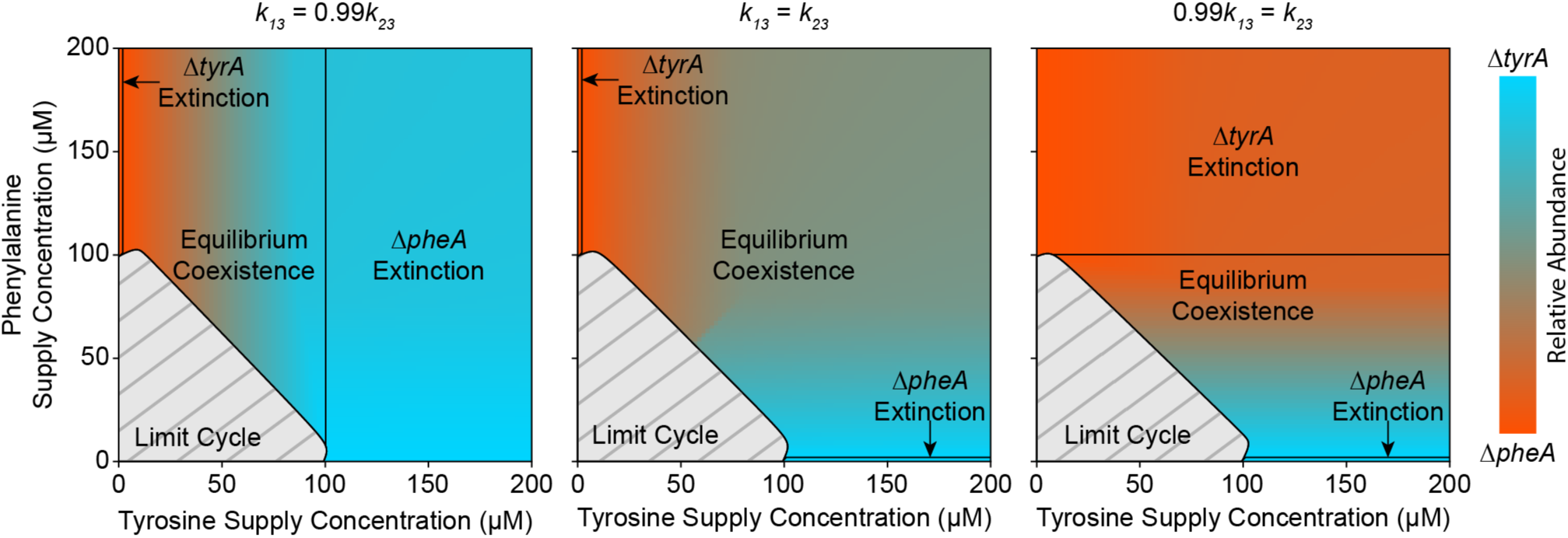
The orientation of transcritical bifurcations and compositional invariance to amino acid depends on the competitive ability of each strain for glucose. Three bifurcation maps are provided based on the full model (equation (1)) using a simplified parameter set (Extended Data Table 1) where all parameters are symmetrical between species except for the glucose half saturation constants *k*_13_ and *k*_23_. Shading between cyan and red maps the steady state community composition to an input of amino acids. Light grey regions with dark grey hashing indicate oscillatory dynamics. Sold black lines represent boundaries between equilibrium coexistence and competitive exclusion. When *k*_13_ is less than *k*_23_ (left plot), then species 1 (*ΔtyrA*) is a better competitor for glucose and the region of *ΔpheA* exclusion is more distinguishing. This community also shows compositional invariance to phenylalanine. The converse is true when *k*_23_ is less than *k*_13_ (right plot). In the perfectly symmetrical case where *k*_13_ is equal to *k*_23_ (middle plot), both amino acids affect the community composition, and the regions of competitive exclusion are minor.

**Extended Data Fig. 4.**
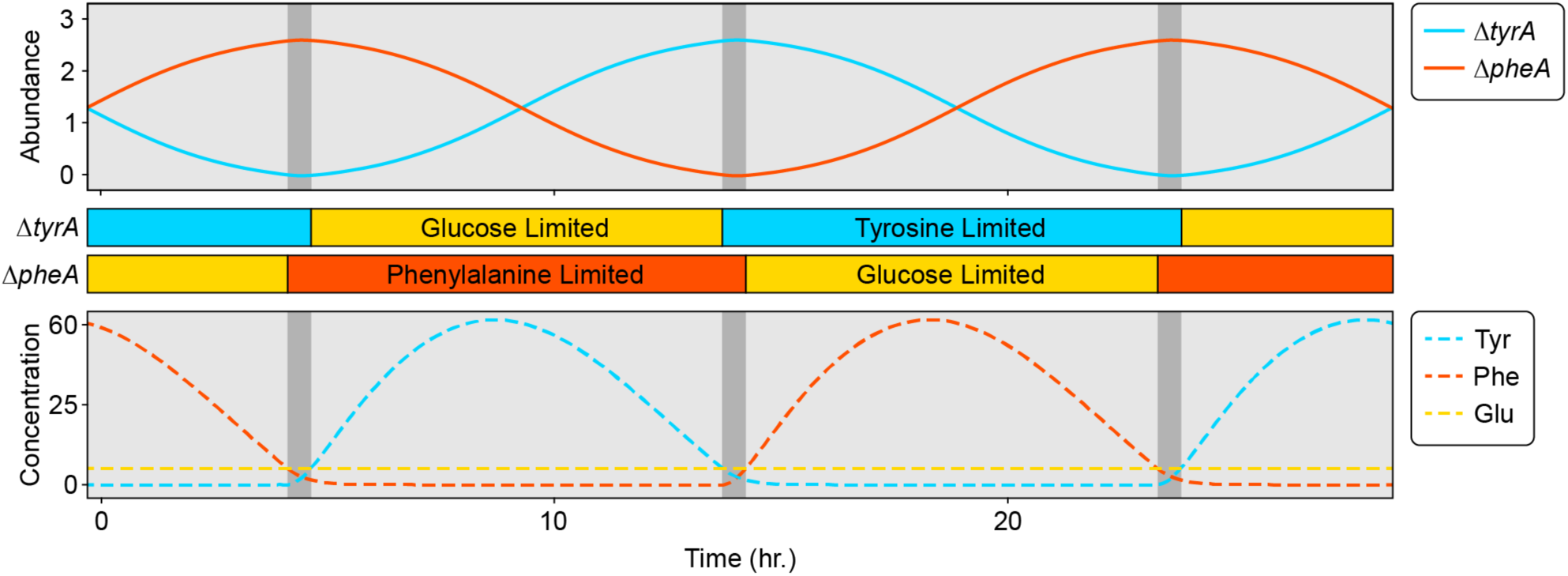
Population oscillations emerge in simulations of the minimal model. Similarly to simulations of the full model (equation (1), Fig. 4a), the auxotroph abundance in the minimal model oscillates between an approximate abundance of 0 and 3, and each cycle occurs over a similar period of time. Resource dynamics show long periods of simultaneous amino acid and glucose limitation (light grey intervals) that are interrupted by short periods of limitation by both amino acids (dark grey intervals). The parameter values of the simulation were: *a*_ij_ = 0.1, *q*_12_ = *q*_21_ = 10, *q*_11_ = *q*_22_ = 20, *R*_3_ = 6.14, *N*_*tot*_ = 2.62, and *D* = 0.2. The initial conditions of the simulation were: *R*_1_ = *R*_2_ = 1, *f* = 0.51.

**Extended Data Fig. 5.**
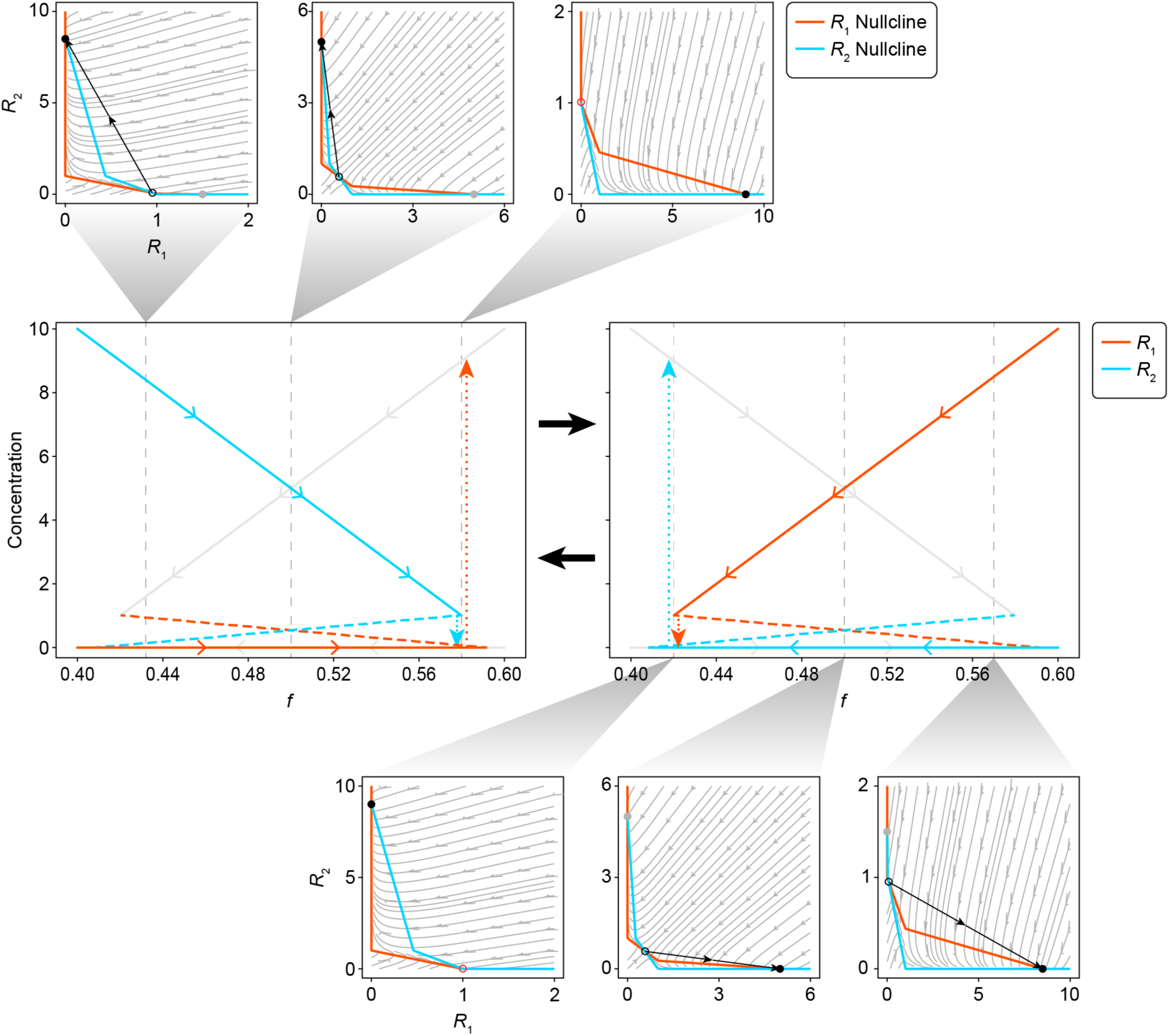
Model reduction leads to a system that exhibits relaxation oscillations when *q*_11_*q*_22_ > *q*_12_*q*_21_is satisfied. The two central plots show the continuation of the *R*_1_ (red) and *R*_2_ (cyan) steady states as a function of fractional abundance in the community. At any given moment, only one of the two sets of steady state solutions are active, which is the distinguishing feature between the two central plots. The inactive steady state solutions are plotted with grey lines. Solid lines are used to indicate stable steady states while dashed lines are used to indicate unstable steady states. Dotted lines indicate the rapid transition between stable states that occurs at the point of annihilation. Each continuation plot is associated with a set of phase portraits showing the *R*_1_ and *R*_2_ streamlines, and how the steady states change with respect to the community composition. The *R*_1_ and *R*_2_ nullclines are plotted in red and cyan respectively. The active stable steady state is shown as a solid black dot, while its inactive counterpart is shown as a solid grey dot. The unstable steady state is shown as an open circle.

**Extended Data Fig. 6.**
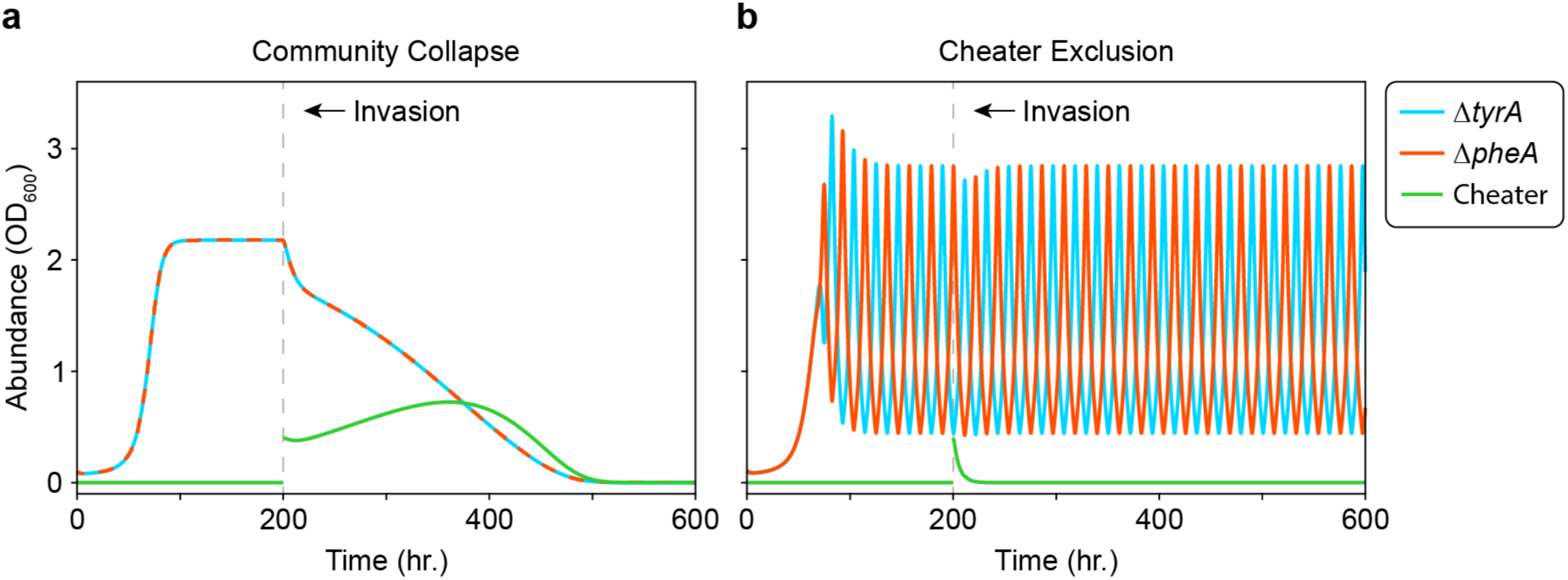
Population cycles from temporal separation of resource production resist invasion and collapse from a metabolic cheater. Each plot shows the dynamics of *ΔtyrA* (cyan), *ΔpheA* (red), and a cheater (green) in a simulated invasion experiment. The cheater was added as an additional variable in the full model (equation (1)) that competes for glucose and amino acid without producing amino acid. Following an initial period with only *ΔtyrA* and *ΔpheA* in the community, the cheater abundance was reset to 0.4 (OD_600_) and the simulation proceeded. A simplified symmetrical parameter set was used for the simulations (Extended Data Table 1), where the resource consumption quotas for the cheater *q*_3j_were equal to those for *ΔtyrA* and *ΔpheA.* Since the cheater was defined as a dual auxotroph, the cheater growth rate was set as the minimum over all three resources with a small growth advantage from the loss of biosynthetic burden where μ_3_ = 1.05 · min(μ_31_, μ_32_, μ_33_). **a**) Invasion of a community at stable equilibrium with the initial condition *N*_1_ = *N*_2_ = 0.1, *N*_3_ = 0, *R*_3_ = 11.11, *R*_1_ = *R*_2_ = 1. **b**) Invasion of a community on its limit cycle with the initial condition *N*_1_ = 0.1, *N*_2_ = 0.11, *N*_3_ = 0, *R*_3_ = 11.11, *R*_1_ = *R*_2_ = 1.

**Extended Data Fig. 7.**
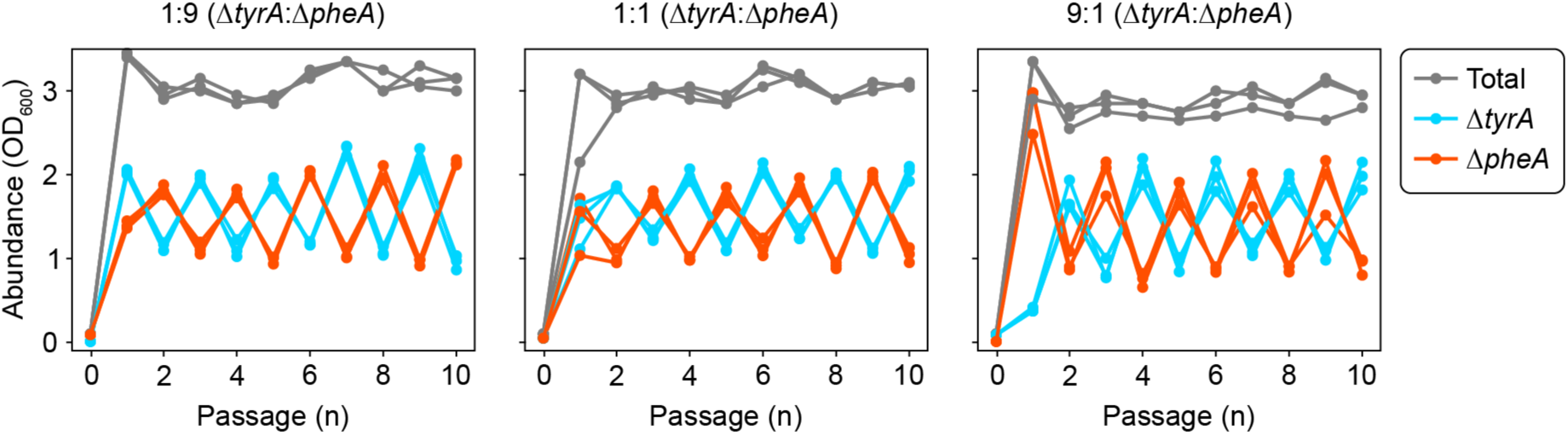
Oscillations are robust to changes in the species inoculum. Communities composed of the *ΔtyrA* and *ΔpheA* auxotrophs (n=3) were established at three different inoculum ratios and passaged for a duration of ten days. One community was inoculated at a *ΔtyrA*:*ΔpheA* ratio of 1:9 (left), another at a ratio of 1:1 (middle), and one more at a ratio of 9:1 (right plot). Each plot shows the absolute abundance of the community members measured at the end of each batch. *ΔtyrA* is plotted in cyan while *ΔpheA* is plotted in red. The total community abundance is plotted in grey. Cultures were established and passaged by diluting the cultures to an initial OD_600_ equal to 0.1 at the beginning of each passage. The media was composed of MOPS Minimal Media (Teknova, #M2106) supplemented with 10 μM tyrosine and 20 μM phenylalanine. While similar oscillations emerged in all inoculum conditions, there was a phase difference between the 1:9 inoculum and the 1:1 and 9:1 inocula. This phase difference occurs at the end of the second batch where the community inoculated at 1:9 was dominated by *ΔpheA* while the other communities were dominated by *ΔtyrA*. The oscillation phase displayed sensitivity to the concentrations of supplemented amino acids when the inoculum was 1:1. This resulted in phase differences between experiments for a single condition dependent on how the amino acid stock solutions were prepared. To improve reproducibility in our presented data, we selected an inoculum ratio of 9:1 for communities of *ΔtyrA* and *ΔpheA*. This always resulted in the same phasing in conditions that produced oscillations.

**Extended Data Table 1.** Parameter values for the full model (Eqn. 1). Each parameter is listed with a brief explanation of its meaning and the values used for all simulations within this work. For parameters that were not assigned a specific value (*R*_1,i*n*_and *R*_2,i*n*_), a range is provided in closed brackets to indicate the sample space of values used for figure generation. The listed parameters were either inferred from a set of experimental data using a nonlinear optimization algorithm in MATLAB (Inferred), or manually chosen to be simple and symmetrical (Simple).

| Parameter | Interpretation | Inferred | Simple |
| --- | --- | --- | --- |
| $\mu_{1,max}$ | Maximum specific growth rate of $\Delta tyrA$ | 0.7045 | 1 |
| $\mu_{2,max}$ | Maximum specific growth rate of $\Delta pheA$ | 0.9517 | 1 |
| $k_{13}$ | Saturation constant for glucose consumption by $\Delta tyrA$ | 3.7551 | 10 |
| $k_{23}$ | Saturation constant for glucose consumption by $\Delta pheA$ | 10.698 | 10 |
| $k_{12}$ | Saturation constant for tyrosine consumption by $\Delta tyrA$ | 20.867 | 10 |
| $k_{21}$ | Saturation constant for phenylalanine consumption by $\Delta pheA$ | 8.6907 | 10 |
| $q_{13}$ | Yield quotient for $\Delta tyrA$ biomass from glucose | 1.2770 | 1 |
| $q_{23}$ | Yield quotient for $\Delta pheA$ biomass from glucose | 3.5704 | 1 |
| $q_{12}$ | Yield quotient for $\Delta tyrA$ biomass from tyrosine | 43.302 | 10 |
| $q_{21}$ | Yield quotient for $\Delta pheA$ biomass from phenylalanine | 39.073 | 10 |
| $q_{11}$ | Maximum production rate of phenylalanine by $\Delta tyrA$ | 80.364 | 20 |
| $q_{22}$ | Maximum production rate of tyrosine by $\Delta pheA$ | 46.602 | 20 |
| $D$ | Dilution rate of the chemostat | 0.2000 | |
| $R_{1,in}$ | Concentration of phenylalanine in the feedstock | [0, 200] | |
| $R_{2,in}$ | Concentration of tyrosine in the feedstock | [0, 200] | |
| $R_{3,in}$ | Concentration of glucose in the feedstock | 11.1111 | |

**Extended Data Table 2.** Best estimate parameter values used for the biomolecular model (Eqn. S1). Each parameter is listed with a brief explanation of its meaning and the value used for all simulations within this work. For parameters that were not assigned a specific value (*R*_1,i*n*_ and *R*_2,i*n*_), a range is provided in closed brackets to indicate the sample space of values used for figure generation. Parameters used for discrete time simulations were inferred from a set of experimental data using a nonlinear optimization algorithm in MATLAB. Parameters specific to the chemostat simulations were manually selected.

| Parameter | Interpretation | Value |
| --- | --- | --- |
| $\mu_{1,max}$ | Maximum specific growth rate of $\Delta tyrA$ | 0.3750 |
| $\mu_{2,max}$ | Maximum specific growth rate of $\Delta pheA$ | 0.8817 |
| $k_{13}$ | Saturation constant for glucose consumption by $\Delta tyrA$ | 1.9620 |
| $k_{23}$ | Saturation constant for glucose consumption by $\Delta pheA$ | 4.5814 |
| $k_{12}$ | Saturation constant for tyrosine consumption by $\Delta tyrA$ | 2.0413 |
| $k_{21}$ | Saturation constant for phenylalanine consumption by $\Delta pheA$ | 6.9279 |
| $q_{13}$ | Yield quotient for $\Delta tyrA$ biomass from glucose | 5.4112 |
| $q_{23}$ | Yield quotient for $\Delta pheA$ biomass from glucose | 4.1842 |
| $q_{12}$ | Yield quotient for $\Delta tyrA$ biomass from tyrosine | 37.7558 |
| $q_{21}$ | Yield quotient for $\Delta pheA$ biomass from phenylalanine | 46.5086 |
| $q_{11}$ | Maximum production rate of phenylalanine by $\Delta tyrA$ | 20.4518 |
| $q_{22}$ | Maximum production rate of tyrosine by $\Delta pheA$ | 11.3321 |
| $k_1$ | Saturation constant for phenylalanine production by $\Delta tyrA$ | 10.7868 |
| $k_2$ | Saturation constant for tyrosine production by $\Delta pheA$ | 2.4787 |
| $n_1$ | Hill coefficient for phenylalanine production by $\Delta tyrA$ | 1.6251 |
| $n_2$ | Hill coefficient for tyrosine production by $\Delta pheA$ | 6.7650 |
| $D$ | Dilution rate of the chemostat | 0.1000 |
| $R_{1,in}$ | Concentration of phenylalanine in the feedstock | [0, 200] |
| $R_{2,in}$ | Concentration of tyrosine in the feedstock | [0, 200] |
| $R_{3,in}$ | Concentration of glucose in the feedstock | 11.1111 |

**Extended Data Table 3.**
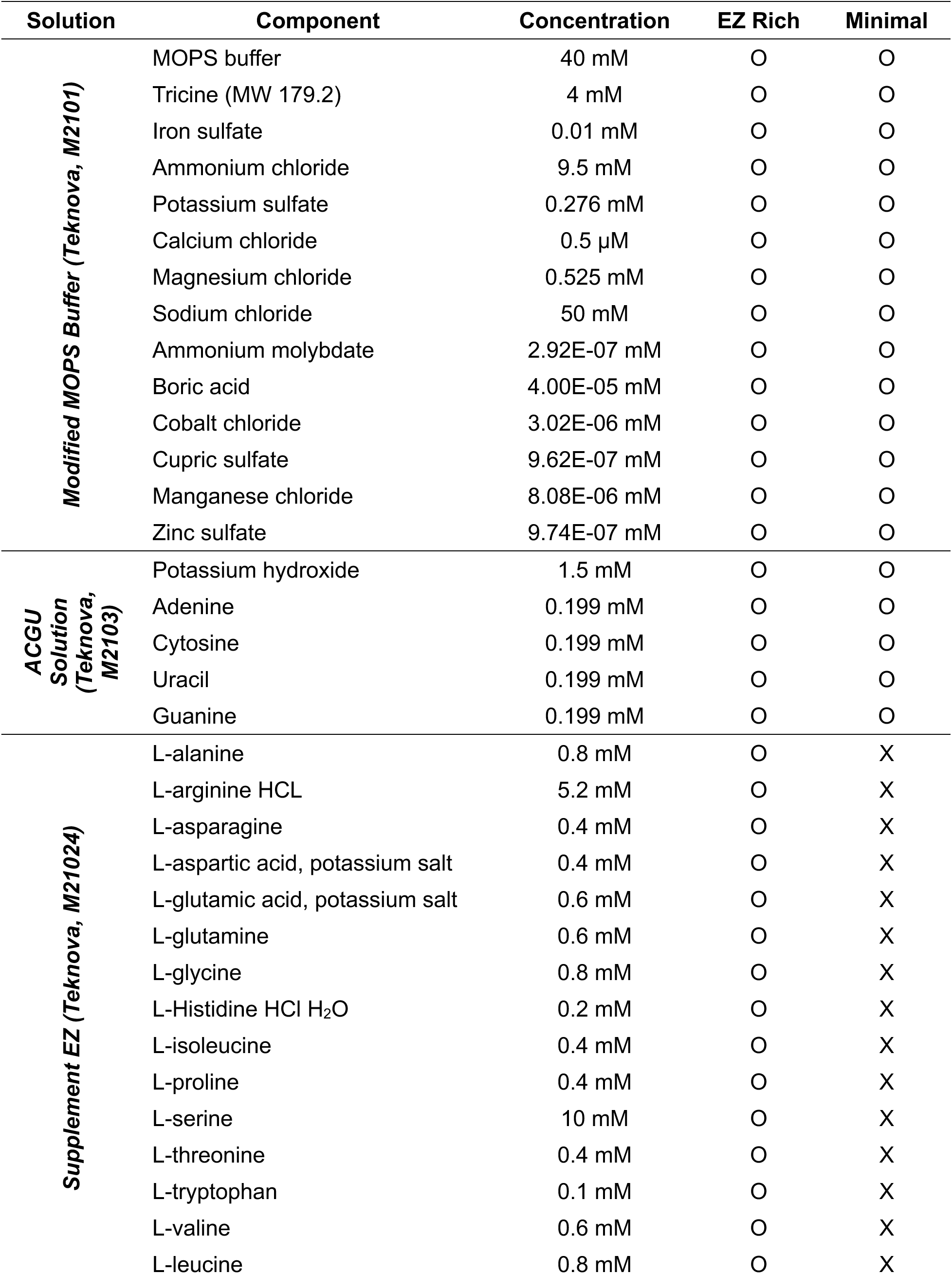
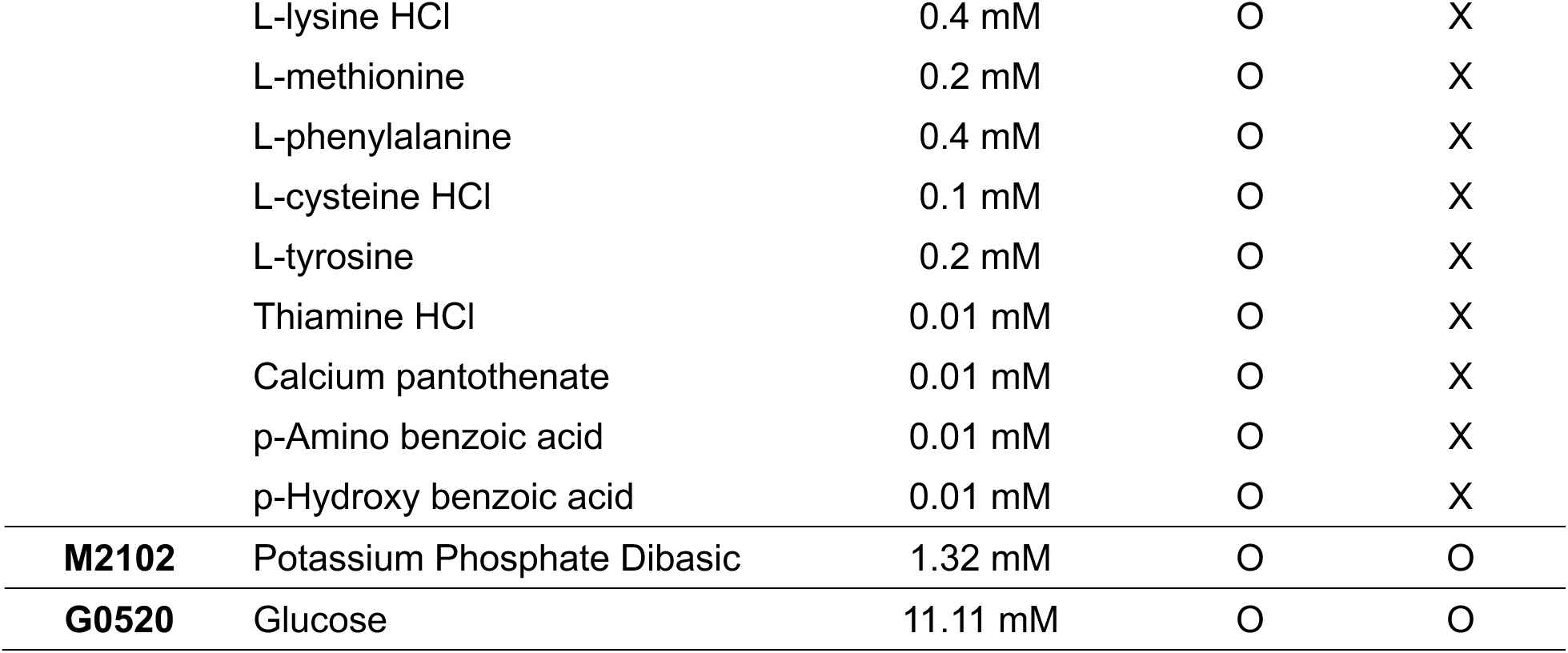
Composition of microbial culture media. Cultures were grown in either EZ Rich Defined Media (Teknova #M2105), MOPS Minimal Media (Teknova, #M2106), or MOPS Minimal Media with various spike in concentrations of tyrosine and/or phenylalanine. All media are chemically defined, and the exact composition is provided here. For the two columns that correspond with either EZ Rich Defined Media or MOPS Minimal Media, the letter O indicates the presence of a specific media component, while the letter X indicates the absence of a component.

| **Solution** | **Component** | **Concentration** | **EZ Rich** | **Minimal** |
| --- | --- | --- | --- | --- |
| **Modified MOPS Buffer (Teknova, M2101)** | MOPS buffer | 40 mM | O | O |
|  | Tricine (MW 179.2) | 4 mM | O | O |
|  | Iron sulfate | 0.01 mM | O | O |
|  | Ammonium chloride | 9.5 mM | O | O |
|  | Potassium sulfate | 0.276 mM | O | O |
| | Calcium chloride | 0.5 μM | O | O |
|  | Magnesium chloride | 0.525 mM | O | O |
|  | Sodium chloride | 50 mM | O | O |
|  | Ammonium molybdate | 2.92E-07 mM | O | O |
|  | Boric acid | 4.00E-05 mM | O | O |
|  | Cobalt chloride | 3.02E-06 mM | O | O |
|  | Cupric sulfate | 9.62E-07 mM | O | O |
|  | Manganese chloride | 8.08E-06 mM | O | O |
|  | Zinc sulfate | 9.74E-07 mM | O | O |
| **ACGU Solution (Teknova, M2103)** | Potassium hydroxide | 1.5 mM | O | O |
|  | Adenine | 0.199 mM | O | O |
|  | Cytosine | 0.199 mM | O | O |
|  | Uracil | 0.199 mM | O | O |
|  | Guanine | 0.199 mM | O | O |
| **Supplement EZ (Teknova, M21024)** | L-alanine | 0.8 mM | O | X |
|  | L-arginine HCL | 5.2 mM | O | X |
|  | L-asparagine | 0.4 mM | O | X |
|  | L-aspartic acid, potassium salt | 0.4 mM | O | X |
|  | L-glutamic acid, potassium salt | 0.6 mM | O | X |
|  | L-glutamine | 0.6 mM | O | X |
|  | L-glycine | 0.8 mM | O | X |
|  | L-Histidine HCl H<sub>2</sub>O | 0.2 mM | O | X |
|  | L-isoleucine | 0.4 mM | O | X |
|  | L-proline | 0.4 mM | O | X |
|  | L-serine | 10 mM | O | X |
|  | L-threonine | 0.4 mM | O | X |
|  | L-tryptophan | 0.1 mM | O | X |
|  | L-valine | 0.6 mM | O | X |
|  | L-leucine | 0.8 mM | O | X |
|  | L-lysine HCl | 0.4 mM | O | X |
|  | L-methionine | 0.2 mM | O | X |
|  | L-phenylalanine | 0.4 mM | O | X |
|  | L-cysteine HCl | 0.1 mM | O | X |
|  | L-tyrosine | 0.2 mM | O | X |
|  | Thiamine HCl | 0.01 mM | O | X |
|  | Calcium pantothenate | 0.01 mM | O | X |
|  | p-Amino benzoic acid | 0.01 mM | O | X |
|  | p-Hydroxy benzoic acid | 0.01 mM | O | X |
| **M2102** | Potassium Phosphate Dibasic | 1.32 mM | O | O |
| **G0520** | Glucose | 11.11 mM | O | O |

## Notes

### Competing Interest Statement

The authors have declared no competing interest.

