## Supplementary Information for "Metabolic interplay drives population cycles in a cross-feeding microbial community"

### Supplementary Discussion

*Auxotrophic escape in amino acid starved cultures of ΔpheA*

The *ΔpheA* auxotroph displayed two distinct periods of growth during a 24-hour batch in phenylalanine limited media (Fig. 2f,g). Growth arrest after the initial period coincided with the depletion of phenylalanine, while growth arrest after the second period coincided with depletion of glucose. Since phenylalanine was depleted during the second period of growth, these results demonstrate the ability of *ΔpheA* to escape auxotrophy between 12 and 24 hours within a batch. Previous studies have demonstrated that this auxotrophic escape can occur in acidic environments where the reaction catalyzed by *pheA* can occur spontaneously[1,2]. Consistent with this notion, the extracellular pH decreased substantially prior to rescue of *ΔpheA* growth between 12 and 24 hours (Fig. S1). This growth rescue phenomenon could be expected to decouple the mutualism in the community context, driving extinction of *ΔtyrA* when amino acids are limiting. However, this was not observed, likely due to the late onset of growth rescue. Thus, it is unlikely that growth rescue has an impact on the dynamical behaviors in our community experiments.

*An alternative model with explicit encoding of resource cross-inhibition according to a biomolecular mechanism captures oscillatory dynamics.*

Our full model (equation (1)) encodes the cross-inhibition of amino acids through a strategy of resource management. Such a general mechanism could be more broadly applicable to interacting microbial communities. To investigate more directly the mechanism that could generate oscillations in this specific community, we considered the metabolic features of our system that could potentially generate cross-inhibition at a biochemical-level. Tyrosine and phenylalanine biosynthesis only diverge at the penultimate reaction[2], and thus share most reaction steps, biosynthetic machinery, intermediates, and regulation strategies. We hypothesize that the gene knockouts *ΔtyrA* and *ΔpheA* lead to dysregulation of the anabolic pathways, leading to emergent cross-inhibitory interactions. In a wild type *E. coli*, tyrosine and phenylalanine production is functional, which results in feedback regulation of the deoxy-D-arabino-heptulosonic acid-7-phosphate (DAHP) synthase isoenzymes *aroG* and *aroF* (Fig. S2a)[3-5]. This limits the buildup of the intermediate chorismate and subsequent production of tyrosine and phenylalanine. Conversely, when either of the bifunctional tyrosine or phenylalanine specific terminal enzymes is removed due to the gene knockouts *ΔtyrA* or *ΔpheA*, the internal concentration of either amino acid could drop significantly lower, relieving negative regulation on one of the DAHP synthases. This might result in a buildup of chorismate which eventually overcomes feedback inhibition (Fig. S2b). Therefore, even in the absence of a direct cross-inhibition mechanism in this regulatory scheme, we observe that the net result appears as if tyrosine is inhibiting phenylalanine production and vice versa.

To determine whether this proposed biomolecular mechanism could generate oscillatory dynamics, we developed an ordinary differential equation model that explicitly encodes the direct cross-inhibitory mechanism of amino acid production (equation (S1)). We fit this model to the experimental data using discrete dilution events described by $D(t) = f\sum_{n=1}\delta(t - n\tau)$, where $f$ is the fraction of culture transferred, $n$ is the passage number, $\tau$ is the period, and $\delta(\cdot)$ is the Dirac delta function. The best parameter estimates are provided in Extended Data Table 2.

$$\frac{dN_1}{dt} = \mu_1 N_1 - DN_1 \quad \text{(S1a)}$$

$$\frac{dN_2}{dt} = \mu_2 N_2 - DN_2 \quad \text{(S1b)}$$

$$\frac{dR_1}{dt} = D\left(R_{1,\text{in}} - R_1\right) + q_{11}\left(\frac{1}{1 + (R_2/k_1)^{n_1}}\right)N_1 - \mu_2 q_{21} N_2 \tag{S1c}$$

$$\frac{dR_2}{dt} = D\left(R_{2,\text{in}} - R_2\right) + q_{22}\left(\frac{1}{1 + (R_1/k_2)^{n_2}}\right)N_2 - \mu_1 q_{12} N_1 \tag{S1d}$$

$$\frac{dR_3}{dt} = D\left(R_{3,in} - R_3\right) - \mu_1 q_{13} N_1 - \mu_2 q_{23} N_2 \tag{S1e}$$

$$\mu_1 = \mu_{1,\max}\left(\frac{R_3}{k_{13} + R_3}\right)\left(\frac{R_2}{k_{12} + R_2}\right) \tag{S1f}$$

$$\mu_2 = \mu_{2,\max}\left(\frac{R_3}{k_{23} + R_3}\right)\left(\frac{R_1}{k_{21} + R_1}\right) \tag{S1g}$$

Simulations of this model accurately recapitulates the stable equilibrium and oscillatory dynamics observed in the 10-batch passaging experiment (Fig. S3a), on which the model was trained. This result supports the notion that a biomolecular mechanism of resource cross-regulation can produce oscillatory dynamics. We investigated whether the periodic forcing through daily passaging was a necessary component for oscillatory dynamics by simulating the system with a constant dilution rate $D$ as in a chemostat. We applied numerical continuation analysis (see Methods) to determine the parameter regime where various dynamical behaviors emerged (Fig. S3b). Similar to our results with the full model, oscillations were present in a parameter regime defined by relatively low concentrations of amino acid inflow. This parameter regime that generates oscillations is contained within a larger regime defined by stable coexistence of both auxotrophs. If the supply concentrations of amino acid are sufficiently large, the model predicts competitive exclusion. In general, this biomolecular mechanistic model produces strikingly similar results to our full model representing a more general strategy of resource management. Notably, a key difference between the models arises where the biomolecular mechanistic model displays bistability of a stable equilibrium and a dynamic equilibrium (oscillations) over a narrow parameter regime. This regime is defined on one side by a Hopf bifurcation and on the other side by a fold bifurcation (Fig. S3c).

*Minimal model*

Our full model (equation (1)) tracks the dynamics of five variables (two microbial strains and three resources). While it can be explored numerically, it is too complicated to yield closed-form analytical results that can be easily understood. Model reduction can aid in model interpretability, elucidating the key interactions needed to recapitulate the oscillatory dynamics and providing theoretical insights. Here, we study a simpler, related model to understand the core interactions and parameter conditions needed for oscillations. To this end, we applied a series of simplifying assumptions to reduce our full model from five to three, and ultimately to two, equations.

First, we assume that the functional response describing the relationship between resources and species growth rate is linear rather than saturating, replacing equation (1g) with $\mu_{ij} = a_{ij}R_j$, where $a_{ij}$ is the affinity of species $i$ for resource $j$. Next, we focus on the dynamics in the absence of an external supply of amino acids, which eliminates the parameters $R_{1,\text{in}}$ and $R_{2,\text{in}}$. Though our full model does not display population cycles in this amino acid supply regime, minor changes to the inferred parameter set are able to recover limit cycle oscillations when amino acids are not supplied externally.

The glucose concentration in the continuous time model only varies by approximately 8% during the period of a fully developed limit cycle in the parameterized model (Fig. 4a). Therefore, we assume that the glucose concentration within the chemostat is constant, eliminating the equation for $dR_3/dt$. Similarly, the total biomass $N_{tot} = N_1 + N_2$ does not vary substantially relative to the abundance of each individual species, so we also assume that $N_{tot}$ is constant. We define a new variable $0 \le f \le 1$ that describes the relative abundance of $N_1$

$$f = \frac{N_1}{N_1 + N_2}$$

whose dynamics can be derived using the quotient rule:

$$\begin{aligned}\frac{df}{dt} &= \frac{(N_1+N_2)\frac{dN_1}{dt} - N_1\left(\frac{dN_1}{dt} + \frac{dN_2}{dt}\right)}{(N_1+N_2)^2} \\ &= \frac{(N_1+N_2)\mu_1 N_1 - N_1(\mu_1 N_1 + \mu_2 N_2)}{(N_1+N_2)^2} \\ &= \frac{(\mu_1 - \mu_2)N_1 N_2}{(N_1+N_2)^2} \\ &= (\mu_1 - \mu_2) f(1-f).\end{aligned}$$

Thus, our minimal model is

$$\frac{dR_1}{dt} = -DR_1 + q_{11}\big(a_{13}R_3 - \min(a_{12}R_2, a_{13}R_3)\big) f N_{tot} - q_{21}\min(a_{21}R_1, a_{23}R_3)(1-f)N_{tot} \quad \text{(S2a)}$$

$$\frac{dR_2}{dt} = -DR_2 + q_{22}\big(a_{23}R_3 - \min(a_{21}R_1, a_{23}R_3)\big)(1-f)N_{tot} - q_{12}\min(a_{12}R_2, a_{13}R_3) f N_{tot} \quad \text{(S2b)}$$

$$\frac{df}{dt} = \big(\min(a_{12}R_2, a_{13}R_3) - \min(a_{21}R_1, a_{23}R_3)\big) f(1-f). \quad \text{(S2c)}$$

Extended Data Fig. 4 shows the dynamics of the minimal model (equation (S2)), which resemble those of the full model (Fig. 4a) for a set of matching parameter values. Thus, the minimal model captures the essence of the mechanism that leads to cycles in the full model.

We also see that the dynamics of the relative abundance $f$ occur on a slower timescale than those of the amino acids $R_1$ and $R_2$, which show rapid changes at turning points of the cycle (Extended Data Fig. 4). This is because the quota parameters $q_{ij}$ times the total biomass $N_{tot}$ are much larger than unity in both this example and our experimentally fit full model (Extended Data Table 1), so that equations (S2a) and (S2b) can change faster than equation (S2c). This separation of timescales suggests a final simplification, where we assume the fast $R_1$-$R_2$ subsystem reaches a quasi-steady state determined by the slow variable $f$. We then examine the dynamics of $f$ on the slower timescale.

The $R_1$-$R_2$ subsystem (Equations (S2a) and (S2b)) has four potential equilibria, depending on which resource limits each species:

1) When $N_1$ is glucose limited and $N_2$ is amino acid limited, $\min(a_{12}R_2, a_{13}R_3) = a_{13}R_3$ and $\min(a_{21}R_1, a_{23}R_3) = a_{21}R_1$, and

$$\hat{R}_1 = 0, \hat{R}_2 = \frac{\big((1-f)a_{23}q_{22} - f a_{13}q_{12}\big)N_{tot}R_3}{D} \quad \text{(S3)}$$

This equilibrium is valid when $a_{12}\hat{R}_2 \le a_{13}R_3$, or equivalently,

$$f \leq \frac{a_{12}a_{23}q_{22}N_{tot} - a_{13}D}{a_{12}(a_{13}q_{12} + a_{23}q_{22})N_{tot}} = f_{\mathrm{crit},1} \tag{S4}$$

It is always stable when valid, based on the negative eigenvalues of the Jacobian matrix

$$J = \begin{bmatrix} -(1-f)a_{21}q_{21}N_{tot} - D & 0 \\ -(1-f)a_{21}q_{22}N_{tot} & -D \end{bmatrix}$$

2) Similarly, when $N_1$ is amino acid limited and $N_2$ is glucose limited, $\min(a_{12}R_2, a_{13}R_3) = a_{12}R_2$ and $\min(a_{21}R_1, a_{23}R_3) = a_{23}R_3$, and

$$\hat{R}_1 = \frac{(fa_{13}q_{11} - (1-f)a_{23}q_{21})N_{tot}R_3}{D}, \hat{R}_2 = 0 \tag{S5}$$

This equilibrium is valid when $a_{21}\hat{R}_1 \leq a_{23}R_3$, or equivalently,

$$f \geq \frac{a_{21}a_{23}q_{21}N_{tot} + a_{23}D}{a_{21}(a_{13}q_{11} + a_{23}q_{21})N_{tot}} = f_{\mathrm{crit},2} \tag{S6}$$

It is always stable when valid, based on the negative eigenvalues of the Jacobian matrix

$$J = \begin{bmatrix} -D & -fa_{12}q_{11}N_{tot} \\ 0 & -fa_{12}q_{12}N_{tot} - D \end{bmatrix}$$

3) When $N_1$ and $N_2$ are both amino acid limited, $\mu_1 = \min(a_{12}R_2, a_{13}R_3) = a_{12}R_2$ and $\mu_2 = \min(a_{21}R_1, a_{23}R_3) = a_{21}R_1$, and

$$\hat{R}_1 = \frac{fq_{11}(a_{13}(D + fa_{12}q_{12}N_{tot}) - (1-f)a_{12}a_{23}q_{22}N_{tot})N_{tot}R_3}{(D + fa_{12}q_{12}N_{tot})(D + (1-f)a_{21}q_{21}N_{tot}) - f(1-f)a_{12}a_{21}q_{11}q_{22}N_{tot}^2} \tag{S7a}$$

$$\hat{R}_2 = \frac{(1-f)q_{22}(a_{23}(D + (1-f)a_{21}q_{21}N_{tot}) - fa_{13}a_{21}q_{11}N_{tot})N_{tot}R_3}{(D + fa_{12}q_{12}N_{tot})(D + (1-f)a_{21}q_{21}N_{tot}) - f(1-f)a_{12}a_{21}q_{11}q_{22}N_{tot}^2} \tag{S7b}$$

This equilibrium is valid when $a_{12}\hat{R}_2 \leq a_{13}R_3$ and $a_{21}\hat{R}_1 \leq a_{23}R_3$, or equivalently, when $f$ lies between $f_{\mathrm{crit},1}$ and $f_{\mathrm{crit},2}$ defined above. We discuss the stability of this equilibrium below.

4) Finally, when $N_1$ and $N_2$ are both glucose limited, $\min(a_{12}R_2, a_{13}R_3) = a_{13}R_3$ and $\min(a_{21}R_1, a_{23}R_3) = a_{23}R_3$, and

$$\hat{R}_1 = -\frac{(1-f)a_{23}q_{21}N_{tot}R_3}{D}, \hat{R}_2 = -\frac{fa_{13}q_{12}N_{tot}R_3}{D} \tag{S8}$$

Since both resources are negative, this potential equilibrium is inconsistent and therefore never valid.

There are two qualitative cases depending on the order of the critical $f$ values:

1) When $f_{crit,1} < f_{crit,2}$, equilibrium 3 is the only valid equilibrium for $f_{crit,1} < f < f_{crit,2}$, and is stable across this range of $f$. It ends at transcritical bifurcations with equilibrium 1 at $f = f_{crit,1}$ and with equilibrium 2 at $f = f_{crit,2}$ (Fig. S4a). In this case, cycles do not occur in the three-equation minimal model (equation (S2)).

2) When $f_{crit,2} < f_{crit,1}$, equilibria 1 and 2 are both valid and stable for $f_{\text{crit},2} < f < f_{\text{crit},1}$. Equilibrium 3 is valid but unstable across this range of $f$, which can be verified by applying the Routh-Hurwitz criteria to the Jacobian matrix. Equilibrium 3 ends at saddle-node bifurcations with equilibrium 1 at $f = f_{crit,1}$ and with equilibrium 2 at $f = f_{crit,2}$ (Fig. S4b). In this case, cycles may occur in the three-equation minimal model (equation (S2)).

How model parameters determine the case can be seen by comparing $f_{\text{crit},1}$ and $f_{\text{crit},2}$. Case 2 occurs when

$$\frac{a_{12}a_{13}a_{21}a_{23}(q_{11}q_{22} - q_{12}q_{21})N_{tot}}{a_{13}^2a_{21}q_{11} + a_{13}a_{23}\,(a_{12}q_{12} + a_{21}q_{21}) + a_{12}q_{23}^2q_{22}} > D \qquad (S9)$$

Since all parameters are non-negative, equation (S9) shows that $q_{11}q_{22} > q_{12}q_{21}$ is a necessary condition for multiple stable equilibria among the resources (and therefore population cycles). Furthermore, increasing the dilution rate $D$ or decreasing the total abundance $N_{tot}$ (related to glucose supply concentration in our full model) make this case less likely.

Having analyzed the fast $R_1$-$R_2$ subsystem, we now investigate the dynamics of the relative abundance $f$ at the various equilibria of the fast subsystem, focusing on case 2 of multiple stable equilibria. At equilibrium 1 (equation (S3)), species 2 cannot grow ($\mu_2 = 0$) because $\hat{R}_1 = 0$, so $df/dt > 0$. Similarly, at equilibrium 2 (equation (S5)), species 1 cannot grow ($\mu_1 = 0$) because $\hat{R}_2 = 0$, so $df/dt < 0$. Thus, the conditions for relaxation oscillations are met. The $R_1$-$R_2$ subsystem equilibrates to one of the stable equilibria. As $f$ slowly changes, the resources to follow this equilibrium until the fold bifurcation is reached, which precipitates a rapid jump to the other stable equilibrium. The direction of change of $f$ reverses and the cycle repeats (Extended Data Fig. 5).

The above analysis assumes a strict separation of time scales between the fast resource dynamics and the slow relative abundance dynamics. To test importance of this assumption, we compared the stability of the minimal model (equation (S2)) determined by the eigenvalues of its Jacobian matrix with the condition for multiple stable equilibria in the fast resource subsystem (case 2 as determined by equation (S9)), along with the simpler, necessary condition $q_{11}q_{22} > q_{12}q_{21}$ (Fig. S5). Overall, the agreement is very close, with only a narrow range of parameters where multiple stable equilibria in the resource subsystem does not lead to cycles in the three-equation minimal model. A Monte Carlo investigation of parameter values found no examples where limit cycles occurred where the resource subsystem did not have multiple stable equilibria. Finally, Fig. 3c shows that the necessary condition $q_{11}q_{22} > q_{12}q_{21}$ is also applicable to the five-equation full model investigated in the main text.

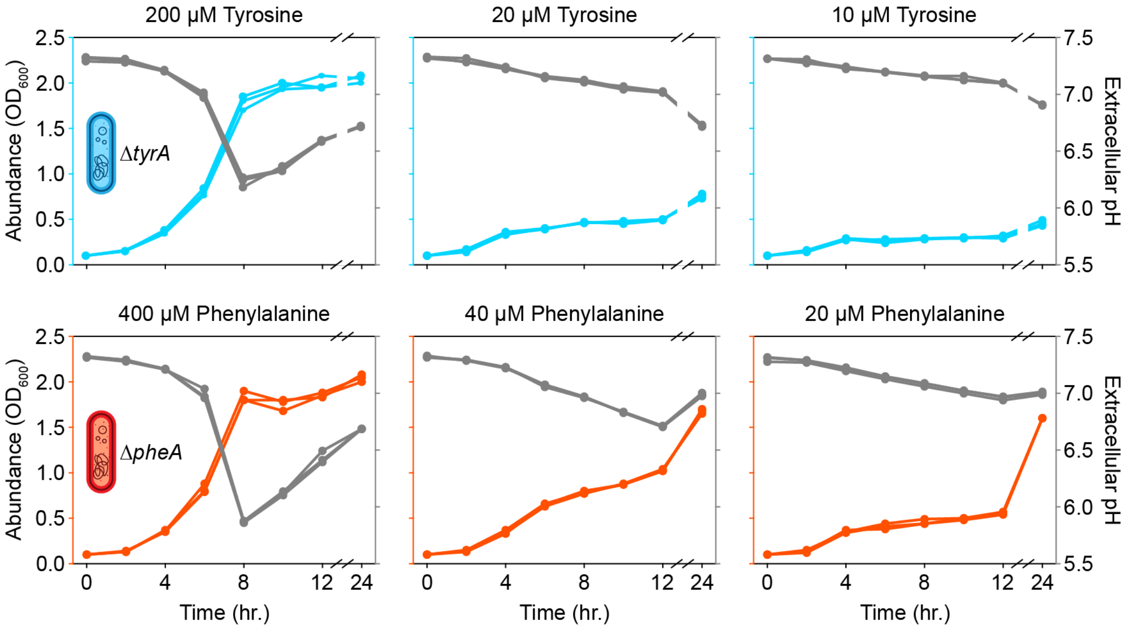


**Fig. S1 | Modification of pH in the extracellular medium by *E. coli*.** Each subplot shows the relationship between growth of a specific amino acid auxotroph and the extracellular pH. The set of subplots is arranged such that each row corresponds with a distinct auxotroph and each column with a distinct media composition. Specifically, the media composition varied in the concentration of each auxotroph's required amino acid. Growth is plotted on the left axis in cyan for *ΔtyrA* and red for *ΔpheA*, and extracellular pH is plotted on the right axis with grey. The x-axis is broken between 12 and 24 hours to condense the plot.

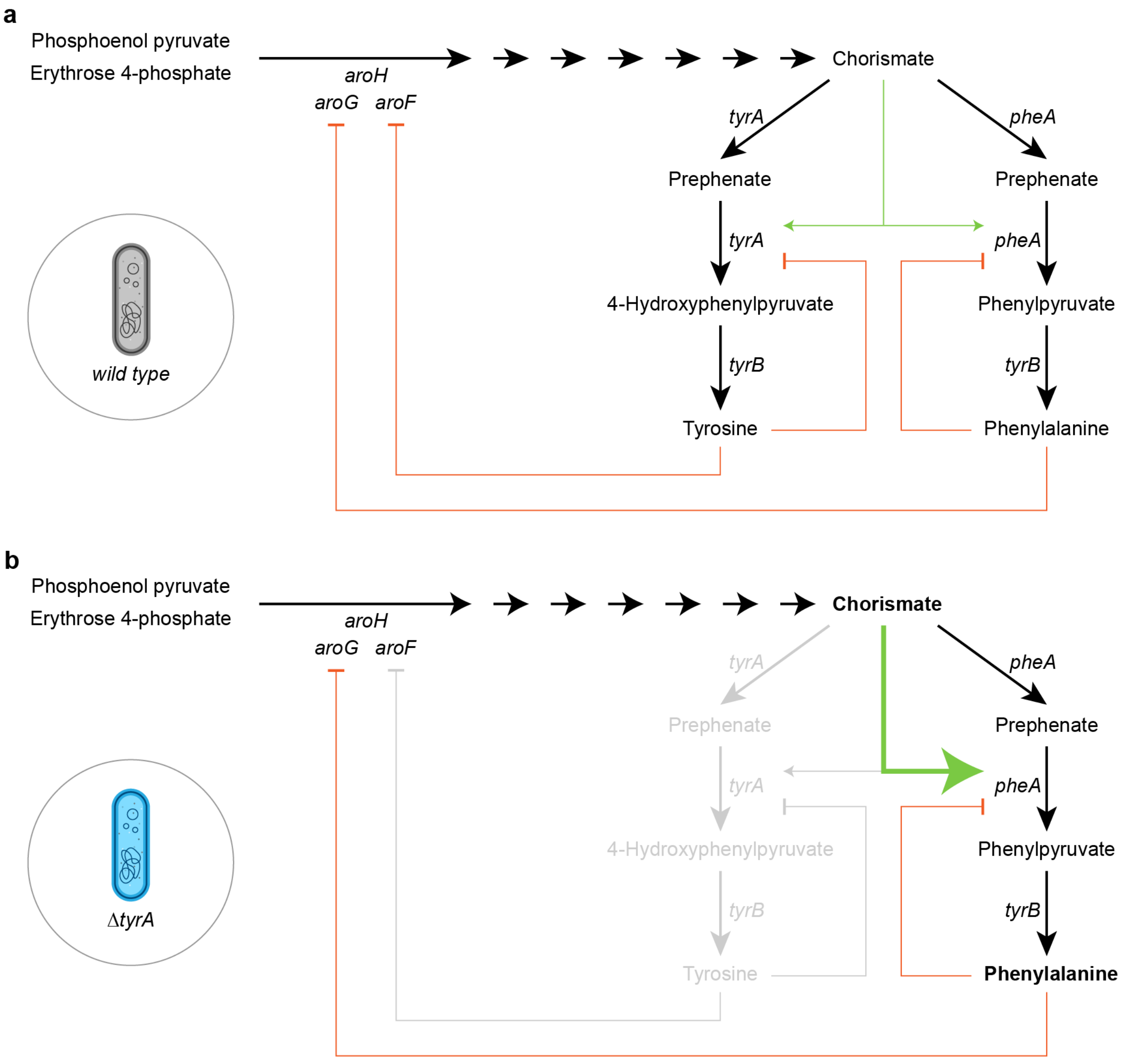


**Fig. S2 | Regulation shifts in the pathway for aromatic amino acid biosynthesis**. Each network diagram highlights the key regulatory steps in the biosynthesis of tyrosine and phenylalanine starting from the central metabolites phosphoenolpyruvate and erythrose 4-phosphate. DAHP synthase isoenzymes catalyze the first committed reaction in the production of chorismate, which is converted into tyrosine and phenylalanine through a series of three sequential reactions. **a**) In wild type *E. coli*, basal production levels of tyrosine and phenylalanine prevent aberrant production of chorismate. **b**) The ability to produce tyrosine is eliminated due to a *ΔtyrA* gene knockout resulting in depletion of intracellular tyrosine during typical growth. The typical response to this depletion is relief of the negative feedback on the *aroF* DAHP synthase in order to increase chorismate production and subsequently tyrosine. However, since the capacity for tyrosine production is eliminated, the pool of chorismate increases without restriction, which fuels the production of phenylalanine.

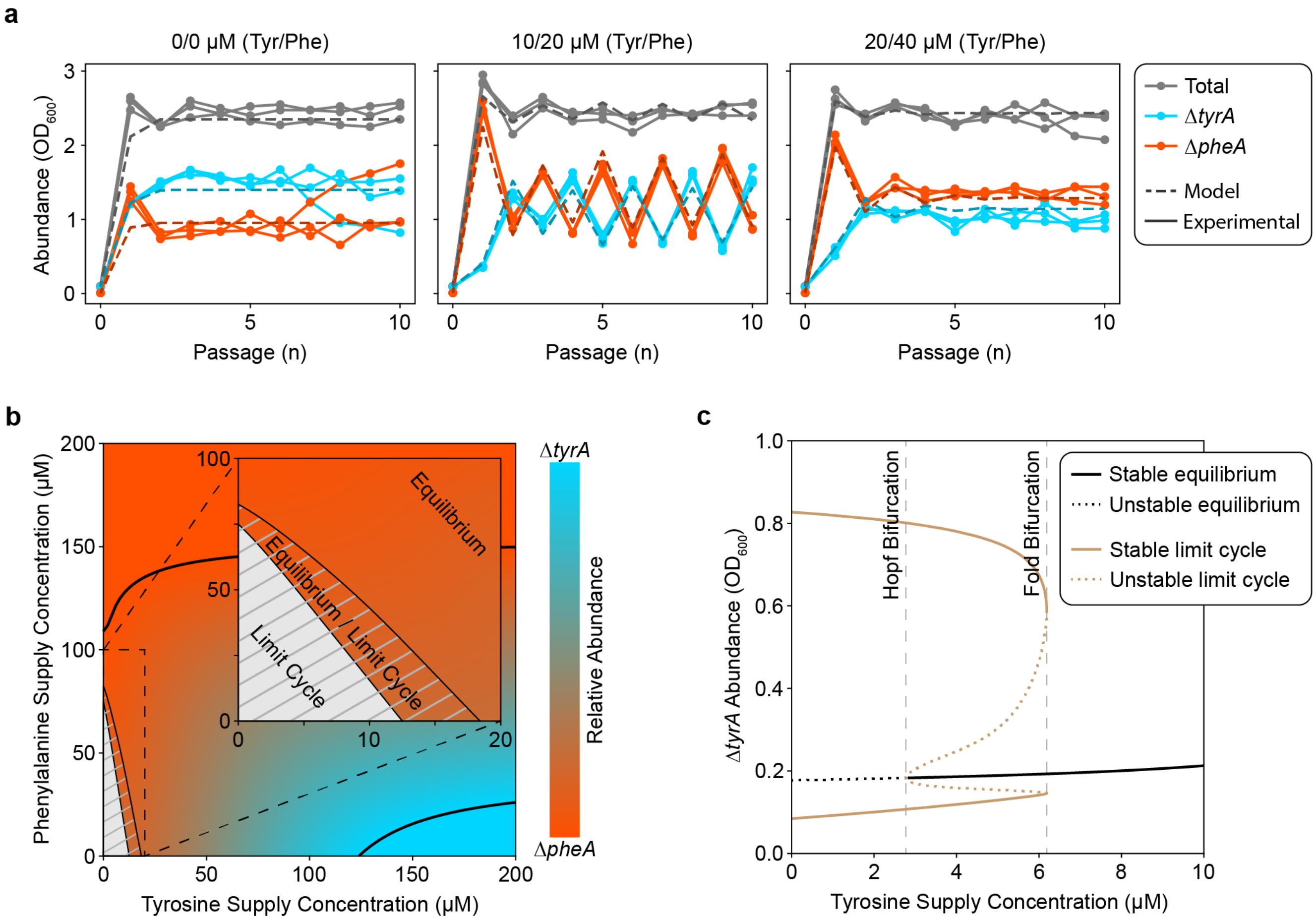


**Fig. S3 | A nonlinear dynamical model representing a biomolecular mechanism of the cross-feeding dynamics recapitulates the stable equilibrium and oscillatory dynamics. a**) Data from the original 10-batch passaging experiment is reproduced with the model performance overlaid with dashed lines. Each plot shows the absolute abundance of community members measured at 24 hours of growth (the duration of a single batch). The amount of amino acid supplementation differentiates between the datasets and is displayed at the top of each subplot. **b**) Numerical continuation analysis and simulations with the continuous time model highlight the parameter regimes where certain dynamical behaviors exist. Both the outset and inset plots show a two-dimensional bifurcation in dynamical behaviors due to changing concentrations of the amino acid inflow. Regions marked with hash lines contain oscillatory dynamics while regions shaded according to the color scale contain a stable equilibrium. Transcritical bifurcations are marked with solid black lines that border regions with saturated colors in which competitive exclusion occurs. The inset plot highlights the regime where only oscillations exist and where bistability exists between stable equilibrium and oscillatory dynamics. **c**) A one-dimensional bifurcation diagram captures changes in fixed point stability through the transition from stable equilibrium only to limit cycle only dynamical behaviors.

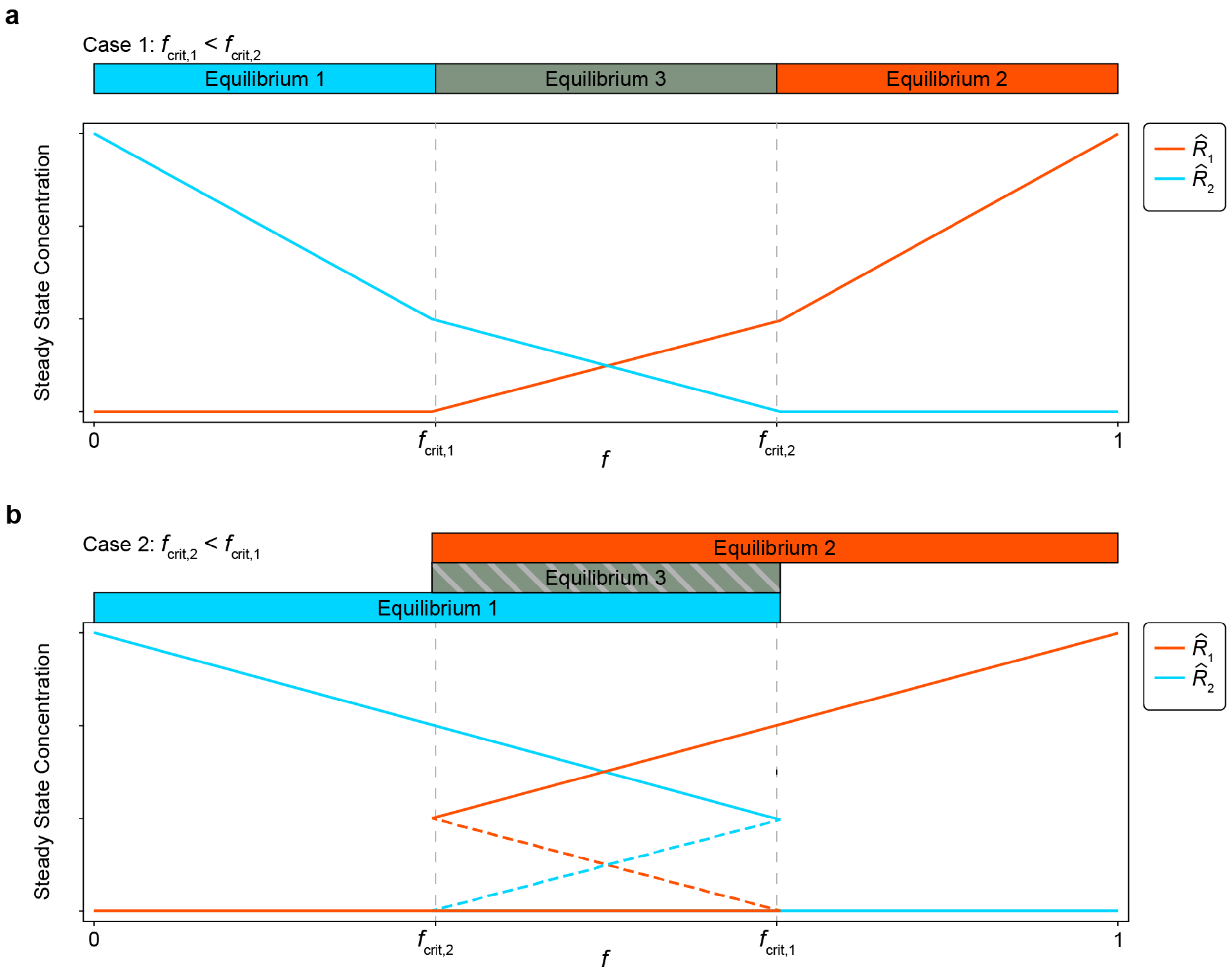


**Fig. S4 | Bifurcation diagrams of the fast $R_1$-$R_2$ subsystem in the minimal model. a**) When $f_{crit,1} < f_{crit,2}$ there is only a single valid, stable equilibrium for every value of $f$. **b**) When $f_{crit,2} < f_{crit,1}$ there are two stable equilibria for $f_{crit,2} < f < f_{crit,1}$ separated by an unstable equilibrium (shown with dashed lines).

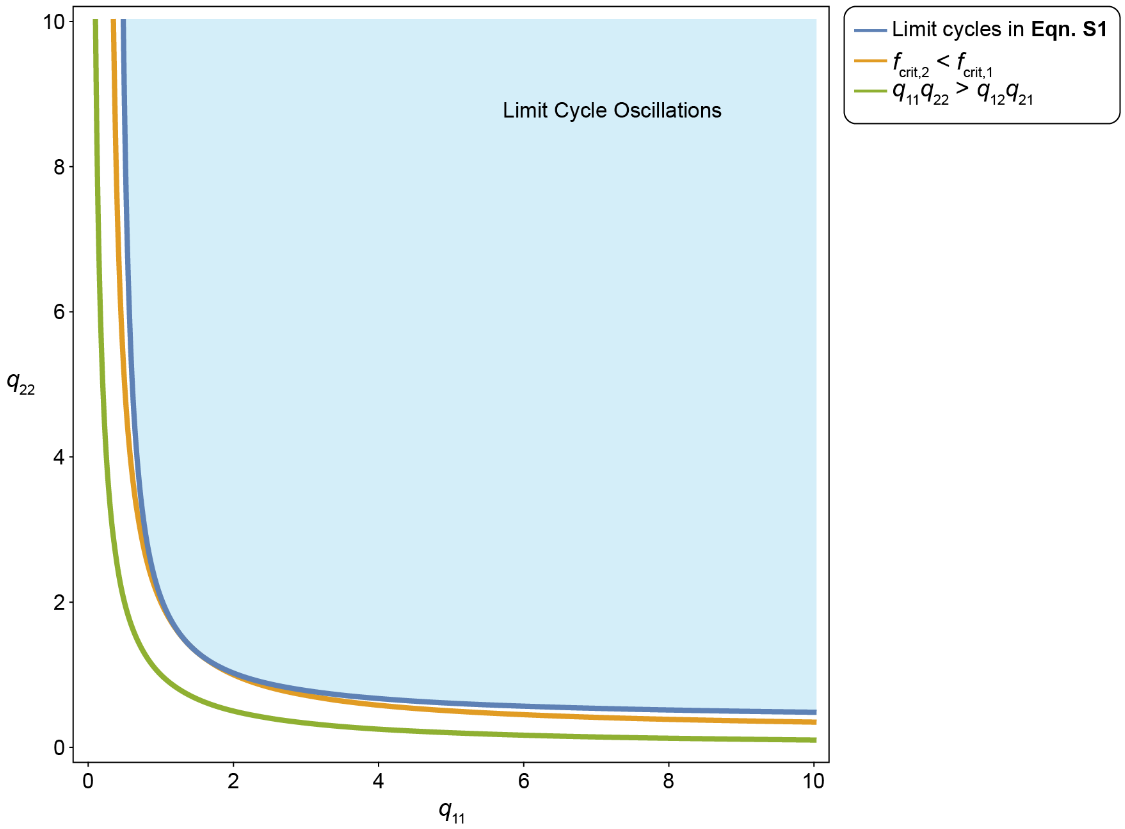


**Fig. S5 | Comparison of three stability criteria in the minimal model**. The blue line represents the stability boundary of the three-equation minimal model (equation (S2)). The yellow line represents the stability boundary of the fast $R_1$-$R_2$ subsystem (equation (S8)). The green line is the necessary condition $q_{11}q_{22} > q_{12}q_{21}$. Parameter values: $N_{tot} = 1$, $R_3 = 1$, $q_{12} = q_{21} = 1$, $a_{12} = a_{21} = 1$, $a_{13} = a_{23} = 1$, $D = 0.2$.
